# Early, adjuvant-responsive epigenetic programs in B cells imprint subsequent plasma cell survival and the duration of humoral immunity

**DOI:** 10.1101/2025.08.15.666880

**Authors:** Tyler J. Ripperger, Lucas J. D’Souza, James F. Read, Wenjie Qi, Darren A. Cusanovich, Stacey Schultz-Cherry, Lynn M. Corcoran, Anthony Bosco, Deepta Bhattacharya

## Abstract

The duration of antibody production varies across different infections and vaccines. To define molecular programs that promote durable humoral immunity, we used mice deficient in ZBTB20, a transcription factor that is highly expressed by plasma cells and required to maintain antibody production *in vivo*. However, genetic deletion of *Zbtb20* in long-lived plasma cells had no impact on the duration of antibody production. Instead, deletion of *Zbtb20* in B cells only within the first week after immunization caused a subsequent failure to maintain plasma cells. Through single-cell ATAC-sequencing, we observed elevated IRF8- and Ets-dependent epigenetic programs in ZBTB20-deficient B cells at 7 days post-immunization. The corresponding transcriptional changes were observed ∼1 week later. Switching from alum to an oil-in-water adjuvant suppressed Ets-dependent epigenetic programs and rescued ZBTB20-deficient antibody responses. Deletion of *Irf8* also rescued ZBTB20-deficient antibody responses. Thus, B cell-intrinsic epigenetic programs imprint durable antibody production at an early stage, prior to major transcriptional consequences and weeks before most long-lived plasma cells are formed.

## INTRODUCTION

The duration of antibody production varies widely across infections and vaccines. For example, vaccines against Human Papilloma Virus (HPV) and Yellow Fever Virus (YFV) induce stable antibody titers that persist for decades^1–7^. In contrast, the seasonal Influenza vaccine and RTS,S Malaria vaccine induce responses that usually wane to background levels within 12 months^8–12^. This durability is primarily a function of the numbers and lifespans of plasma cells, which are terminally differentiated cells downstream of B lymphocytes that secrete enormous quantities of antibodies. In a canonical T cell-dependent antibody response, a large number of plasma cells form within the first few weeks of the response. Most early plasma cells are localized in the extrafollicular regions of secondary lymphoid organs and tend to be short-lived, persisting for only a few days to several weeks^13–16^. As the response progresses, the proportion of plasma cells that live for longer periods increases^17–19^. These longer-lived plasma cells often exit secondary lymphoid organs and traffic to the bone marrow where they can persist and secrete antibodies for longer periods of time^20^, sometimes even for life^21,22^.

Some aspects of the long-lived plasma cell programs are induced in the bone marrow after their formation, where signals such as CD80/86, APRIL, and extracellular ATP promote survival^23–26^. Yet substantial evidence also suggests that key aspects of the plasma cell survival program are imprinted early in the response^27–30^, prior to their migration to the bone marrow. For example, antigens with high valency and assembled into virus-like particles tend to generate more durable antibody titers than antigens that are monomeric^31,32^. Because most plasma cells have dampened surface B cell receptor expression and signaling capacity^33^, these multimeric antigens likely act on upstream B cells. Similarly, specific innate triggers and adjuvants can influence the duration of antibody production^34–37^. Some of these adjuvants may prolong germinal center (GC) reactions and the window of plasma cell formation^38,39^, but much of their activity occurs very early in the response on antigen-presenting cells and potentially directly on B cells^40–42^. Early inflammatory signals clearly influence the subsets of memory B cells that are generated^43^. Yet despite the circumstantial evidence implicating the importance of the early imprinting of longevity, the molecular identity of such programs is not known for plasma cells. Therefore, the concept remains theoretical. Defining a core set of molecular correlates and functionally important features, if measured and replicated across vaccines, could provide a template to generate durable immunity irrespective of the pathogen. Such efforts have implicated platelet and megakaryocyte signatures as extrinsic correlates of durable immunity^26^, but no analogous B cell- and/or plasma cell-intrinsic programs have yet been identified.

We previously identified the transcription factor ZBTB20 as highly expressed by plasma cells and antigen-experienced B cells^36,44,45^. ZBTB20 belongs to a family of factors characterized by an N-terminal BTB-POZ domain that mediates homodimerization and recruitment of transcriptional repressors, and a variable number of zinc finger domains in the C-terminus that mediate sequence-specific DNA binding^46,47^. Following alum-adjuvanted immunizations, *Zbtb20* gene-trapped or knockout fetal liver chimeras failed to sustain long-lived antigen-specific plasma cells^36,44^. Mutant chimeras showed normal GC numbers and organization, affinity maturation, and plasma cell and memory B cell formation. Therefore, our interpretation was that ZBTB20 likely acts directly in plasma cells to promote survival. Yet the germline nature of the *Zbtb20* mutations and the infrequency of plasma cells in mutant chimeras precluded experiments from testing this interpretation directly and defining molecular mechanisms. Here, using inducible knockout mice and single-cell profiling, we instead found evidence of an early requirement for ZBTB20 and a B cell-intrinsic epigenetic imprint that promotes durable antibody responses. This imprint is evident within the first week after immunization when transcriptional consequences are subtle, and weeks before most long-lived plasma cells are formed. Such programs might potentially be used as a target for vaccine engineering and considerably reduce the timeframe needed to iteratively optimize the durability of immunity.

## RESULTS

### ZBTB20 deficiency in B cell lineages induces impaired antibody responses following alum-adjuvanted immunizations

Using a transcriptional reporter, we had previously shown that ZBTB20 expression is minimal in resting naïve B cells but elevated in GC B cells, memory B cells, and plasma cell lineages^36,44^. Using intracellular flow cytometric staining of ZBTB20 protein, we confirmed this expression pattern in antigen-specific cells of mice immunized with 4-hydroxy-3-nitrophenyl acetyl-chicken gamma-globulin (NP-CGG) precipitated in alum adjuvant (**Figures 1A and S1A**). We also observed ZBTB20 protein expression in GL7^positive^ CD38^positive^ antigen-specific pre-GC activated B cells^48^ (B blasts) **(Figures 1A and S1A)**.

**Figure 1:**
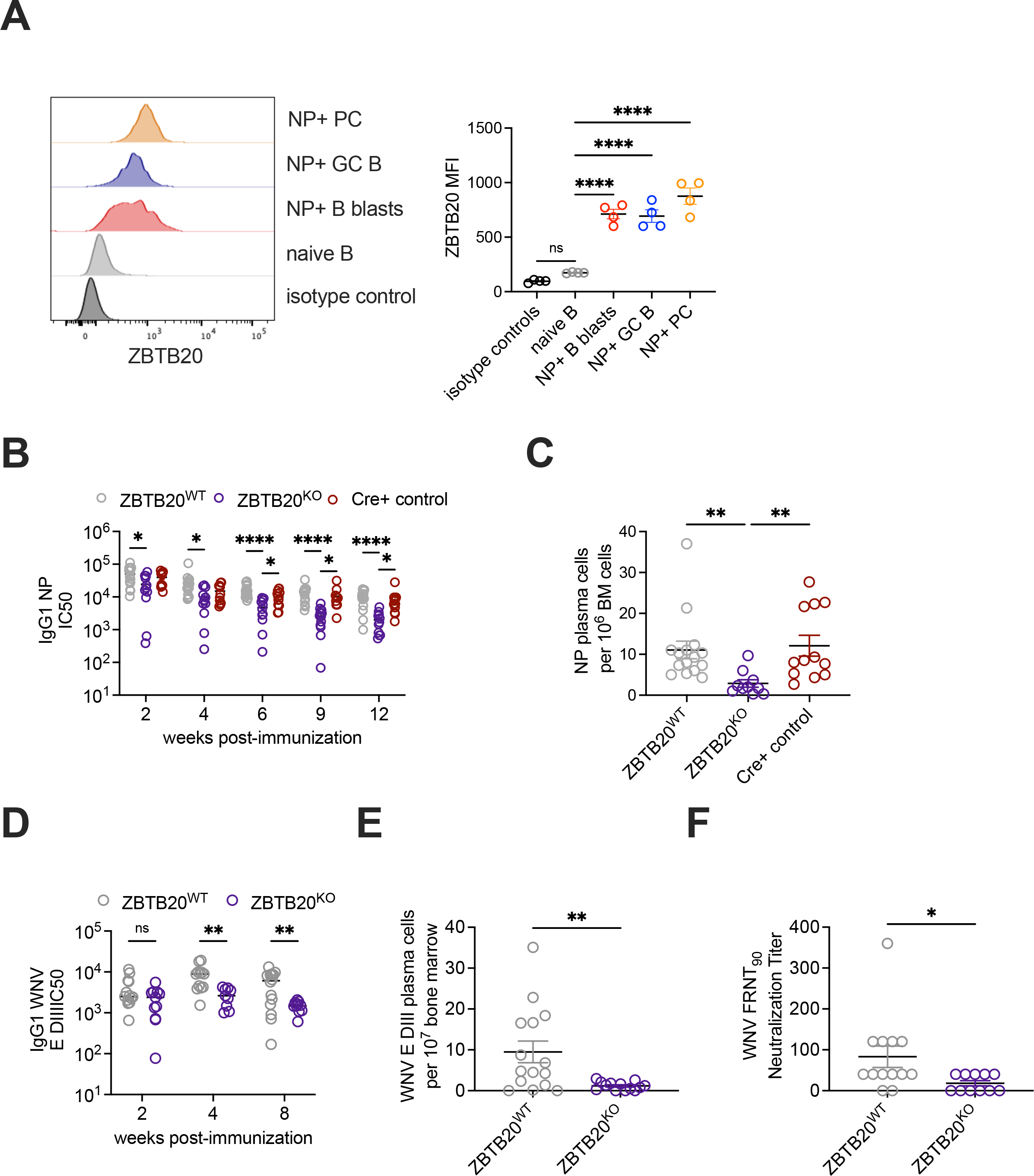
ZBTB20 deficiency in B cells impairs the durability of antibodyresponses. (**A**) ZBTB20 protein expression was assessed by intracellular flow cytometry staining across B cell subtypes in ZBTB20^WT^ mice 7 days post-immunization of NP-CGG/alum. PC=plasma cells; GC=germinal center. Representative histograms of ZBTB20 expression from individual mice across B cell subtypes are shown (left) with quantification of mean fluorescence intensities (MFI) of ZBTB20 staining (right). Each dot represents an individual mouse. Data are representative of three individual experiments. Mean values ± SEM are shown. *= p-value <0.05 ** = p-value < 0.01, *** = p-value < 0.001, **** = p-value < 0.0001 by ordinary one-way ANOVA with Tukey multiple comparisons test. (**B**) IgG1 anti-NP serum antibodies were quantified following NP-CGG/alum immunization via ELISA in ZBTB20^KO^, ZBTB20^WT^, and age-matched Cre+ controls. The IC50 values for individual mice were calculated from serial dilution curves at each time point indicated. Each dot represents an individual mouse. Cumulative data from three individual experiments is shown. Mean values ± SEM are shown. *= p-value <0.05, ** = p-value < 0.01, *** = p-value < 0.001, **** = p-value < 0.0001, by two-way ANOVA with mixed-effects model and Tukey multiple comparisons testing for IC50 values of each genotype across indicated time points. (**C**) IgG1 NP specific bone marrow plasma cells were quantified via ELISPOT at 12 weeks post-immunization in ZBTB20^KO^, ZBTB20^WT^, and age-matched Cre+ controls. Each dot represents an individual mouse. Cumulative data from three individual experiments is shown. Mean values ± SEM are shown. **= p-value <0.01 by Kruskal-Wallis test with Dunn’s multiple comparisons. (**D**-**F**) ZBTB20^KO^ mice and ZBTB20^WT^ littermate controls were immunized with WNV E-DIII-ODN1826 in alum and antibody responses were assessed. (**D**) IgG1 anti-E-DIII serum antibody titers were quantified longitudinally by ELISA. Each dot represents an individual mouse. Cumulative data from two individual experiments is shown. Mean values ± SEM are shown. * = p-value <0.05, ** = p-value < 0.01, by two-way ANOVA with mixed-effects model with Šídák multiple comparison of IC50 values of each genotype across indicated time points. (**E**) IgG1 WNV DIII-specific bone marrow plasma cells were quantified via ELISPOTs in ZBTB20^KO^ mice and ZBTB20^WT^ at 8 weeks post-immunization. Each dot represents an individual mouse. Cumulative data from two individual experiments is shown. Mean values ± SEM are shown. * = p-value <0.05, ** = p-value < 0.01 by unpaired Welch’s two-tailed T-test. (**F**) In-vitro focus reduction neutralizing activity against WNV (NY99) of serum from ZBTB20^KO^ mice and ZBTB20^WT^ littermate controls was assessed 4 weeks post-immunization. Each dot represents an individual mouse. Cumulative data from two individual experiments is shown. Mean values ± SEM are shown. * = p-value <0.05 by unpaired Welch’s two-tailed t-test.

To confirm the defects in durable antibody production we previously observed in gene-trapped and knockout fetal liver chimeras, we crossed *Zbtb20^fl/fl^* mice to the *Mb1-Cre* driver strain to achieve B cell-specific deletion^49,50^. As ZBTB20 expression is minimal in naïve B cells, we validated ZBTB20 protein loss in *Zbtb20^fl/fl^ Mb1-Cre*+ mice in antigen-specific GC B cells compared to littermate *Zbtb20^fl/fl^* controls **(Figure S2A)**. RNA-Seq analysis of GC B cells also demonstrated elimination of transcripts within the floxed exon in *Zbtb20^fl/fl^ Mb1-Cre*+ mice **(Figure S2B**). For simplicity, we refer to *Zbtb20^fl/fl^ Mb1-Cre+* mice as ZBTB20^KO^. ZBTB20^KO^ mice were immunized with NP-CGG/alum alongside *Zbtb20^fl/fl^ Mb1-Cre-* (ZBTB20^WT^) littermate controls and age-matched *Zbtb20^+/+^ Mb1-Cre+* controls. NP-specific serum antibody responses were measured by ELISA for 12 weeks post-immunization. Antibody titers were similar in ZBTB20^KO^ mice relative to controls at 2 weeks post-immunization, yet defects became apparent over time **(Figure 1B)**, consistent with our previous observations^36,44^. ELISPOT analysis at 12 weeks post-immunization confirmed a reduction of antigen-specific bone marrow plasma cells in ZBTB20^KO^ mice **(Figure 1C)**. Splenic plasma cells in ZBTB20^KO^ mice were not statistically significantly reduced relative to ZBTB20^WT^ littermate controls (p=0.06, **Figure S2C**), whereas a deficiency in serum NP-specific IgM was observed at 2 weeks post-immunization **(Figure S2D)**. To confirm that these defects extended to non-haptenated immunogens, domain III (D-III) of the WNV-envelope (E) protein was mixed with the murine TLR9 ligand ODN 1826, precipitated with alum adjuvant, and used to immunize mice. WNV E-DIII displays strong immunogenicity and is an immunodominant target of neutralizing antibodies in mice^51–53^. WNV E-DIII specific antibodies in ZBTB20^KO^ mice decreased over time relative to control animals, and antigen-specific plasma cells in the bone marrow were also diminished at the conclusion of the experiment **(Figure 1D-E)**. Moreover, sera from ZBTB20^KO^ mice were impaired in their ability to neutralize WNV *in vitro* relative to controls **(Figure 1F).** Collectively, these findings confirm our prior observations in *Zbtb20* gene-trapped chimeras using an independent strain and suggest that TLR ligands do not rescue the antibody defect. To define the B cell stages at which ZBTB20 is functionally important, we next employed tamoxifen-inducible systems for deletion.

#### ZBTB20 promotes plasma cell durability upstream of long-lived plasma cell differentiation

ZBTB20 promotes the longevity of plasma cell responses and is highly expressed by bone marrow plasma cells^36,44^. Moreover, overexpression of ZBTB20 in culture induces expression of plasma-cell transcription factors and pro-survival genes^44^. Accordingly, we hypothesized that ZBTB20 function was essential in the long-lived plasma cell stage. To test this hypothesis, we crossed *Zbtb20^fl/fl^* animals with *J-Chain IRES CreERT2* mice, which express tamoxifen-inducible Cre recombinase in plasma cells^52^. After oral gavage of tamoxifen, *Zbtb20^fl/fl^ J-ChainCreERT2^+/^*^-^ (ZBTB20^PC KO^) lost ZBTB20 protein in bone marrow plasma cells **(Figure S3A)**. We reasoned that this system would allow us to capture and transcriptionally profile ZBTB20-deficient plasma cells before they die to define critical survival programs.

We immunized ZBTB20^PC KO^ mice and littermate and age-matched *J-ChainCreERT2+* controls with NP-CGG/alum. Mice were then left untreated until 8 weeks post-immunization to allow for generation of the long-lived plasma cell compartment^17,54^. At 8 weeks, we induced ZBTB20 deletion with tamoxifen and tracked the subsequent antigen-specific antibody titers. Contrary to our hypothesis, we observed no decline in NP-specific antibody titers in ZBTB20^PC KO^ mice relative to controls **(Figure 2A)**. We also observed no differences in antigen-specific bone marrow plasma cell numbers between ZBTB20^PC KO^ mice and controls at 14 weeks as measured by ELISPOT **(Figure 2B).** These results suggested that ZBTB20 is dispensable for plasma cell maintenance once this compartment has been formed. Instead, ZBTB20 may act at the time of or before long-lived plasma cell formation.

**Figure 2:**
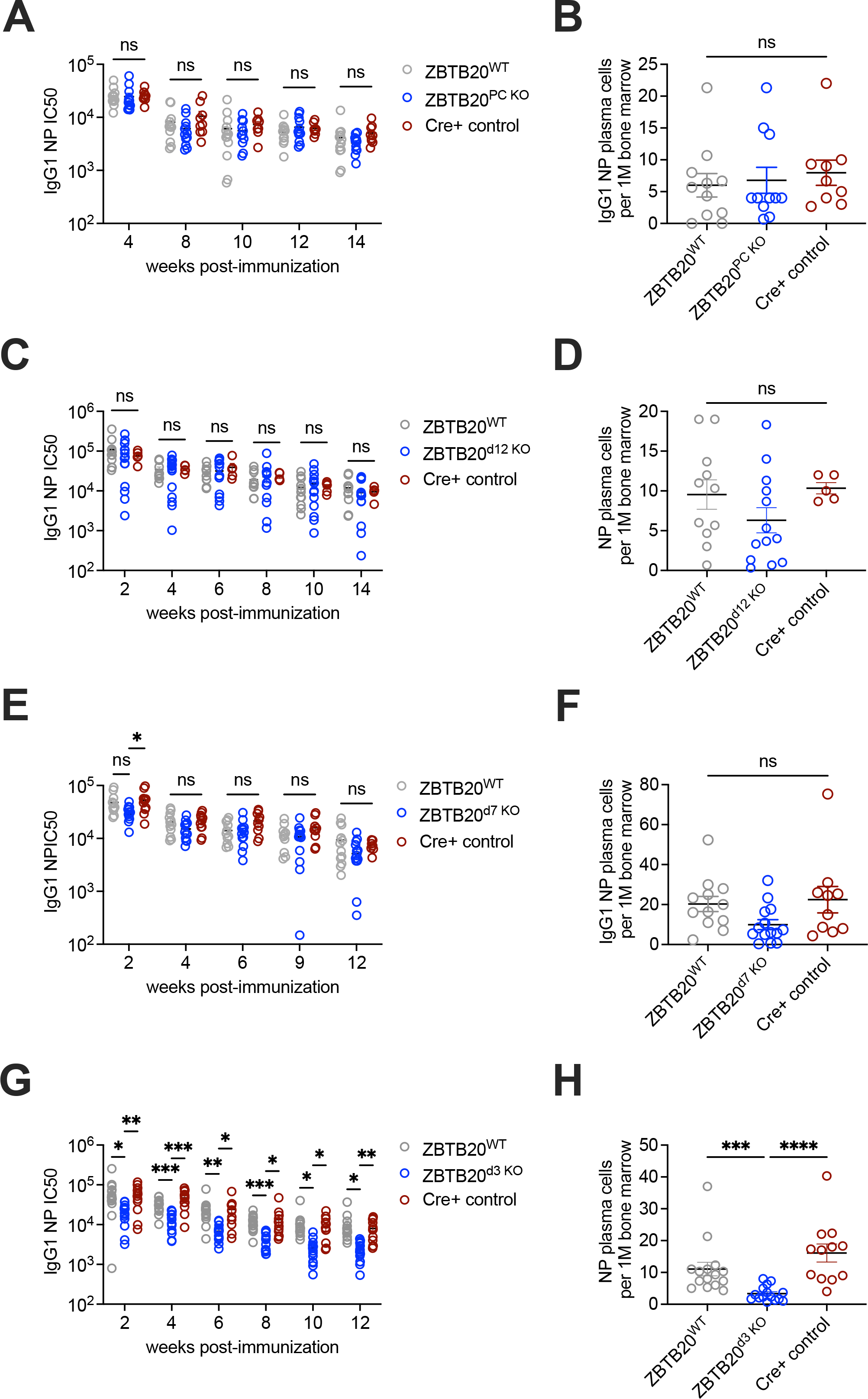
ZBTB20 promotes durable antibody production *in vivo* prior to long-lived plasma cell formation. **(A-B)** ZBTB20^PC KO^ (blue), littermate ZBTB20^WT^ controls (grey), and age-matched Cre+ controls (red) were immunized with NP-CGG/alum. Tamoxifen was administered 8 weeks post-immunization. (A) IgG1 NP specific serum antibodies were assessed longitudinally. Each dot represents an individual mouse. Data are cumulative of two individual experiments. Mean values ± SEM are shown. No statistical significance observed by two-way ANOVA with mixed-effects model and Tukey multiple comparisons testing for IC50 values of each genotype across indicated time points. (B) IgG1 NP specific plasma cells were quantified via bone marrow ELISPOTs at 14 weeks post-immunization in ZBTB20^PC KO^, littermate ZBTB20^WT^ controls, and age-matched Cre+ controls. Each dot represents an individual mouse. Data are cumulative of two individual experiments. Mean values ± SEM are shown. Statistical significance by Kruskal-Wallis test with Dunn’s multiple comparisons. (**C-H**) *Zbtb20^fl/fl^* mice were crossed to hCD20 CreERT2 mice to generate inducible depletion of ZBTB20 in the mature B cells but not differentiated plasma cells. Cre-mediated depletion of ZBTB20 was initiated post- NP-CGG/alum immunization at 12 days (C,D) 7 days (E-F) or 3 days (G-H). In (C,E,G), IgG1 anti-NP specific serum antibodies were quantified via ELISAs longitudinally. Each dot represents an individual mouse. Data are representative of two cumulative experiments for each timed deletion condition. Mean values ± SEM are shown. * = p-value <0.05, ** = p-value < 0.01, *** = p-value < 0.001, **** = p-value < 0.0001 by two-way ANOVA with mixed-effects model and Tukey multiple comparisons testing of IC50 values for each genotype across indicated time points. In (D,F,H) IgG1 NP specific bone marrow plasma cells were quantified via ELISPOT at 14 weeks post-immunization in (D) and 12 weeks post-immunization in (F,H). Each dot represents an individual mouse. Data are cumulative of two individual experiments for each timed deletion condition. Mean values ± SEM are shown. *= p-value <0.05, ** = p-value < 0.01, *** = p-value < 0.001, **** = p-value < 0.0001 by Kruskal-Wallis test with Dunn’s multiple comparisons.

We therefore crossed *Zbtb20^fl/fl^* mice to transgenic *hCD20 CreERT2* animals^55^. This Cre system permits temporal deletion of floxed loci exclusively in B cells, including GC B cells, but not long-lived plasma cells. Following tamoxifen gavage of *Zbtb20^fl/fl^* hCD*20 CreERT2* mice, ZBTB20 protein was efficiently lost in antigen-specific GC B cells **(Figure S3B)**. Tamoxifen treatment of *Zbtb20^fl/fl^ hCD20 CreERT2* mice prior to immunization with NP-CGG/alum led to a defect in antibody responses over time and fewer antigen-specific plasma cells as measured by ELISPOT **(Figures S3C-D),** similar to the ZBTB20^KO^ phenotype.

In the NP-CGG/alum immunization system, most long-lived plasma cells are formed after 2 weeks post-immunization and mostly from the GC^17,18^. To delete *Zbtb20* before most long-lived plasma cells are formed, we administered oral tamoxifen 12 days post-immunization to *Zbtb20^fl/fl^* hCD*20 CreERT2* mice (ZBTB20^d^^12^ ^KO^). ZBTB20^d^^12^ ^KO^ mice demonstrated no defect in their ability to maintain NP-specific serum responses for at least 14 weeks or in the number of bone marrow plasma cells at the conclusion of the experiment **(Figures 2C-D)**. Similar results were observed when tamoxifen was administered at 7 days post-immunization **(Figures 2E-F)**. ZBTB20 protein loss was confirmed via flow cytometry in GC B cells within 3 days following tamoxifen gavage **(Figure S3E)**.

We next administered tamoxifen to *Zbtb20^fl/fl^ hCD20 CreERT2+* mice and controls at 3 days post-immunization. At this timepoint, similar to the results from ZBTB20^KO^ mice, we observed a decline in antibody titers over time in ZBTB20^d^^3^ ^KO^ mice compared to littermate Cre- and age-matched Cre+ controls **(Figure 2G)**. We also observed a reduction in the number of antigen-specific plasma cells in the bone marrow of ZBTB20^d^^3^ ^KO^ animals at 12 weeks post-immunization **(Figure 2H)**. These results indicate that ZBTB20 imprints plasma cell longevity in a narrow window early post-immunization, well before most long-lived plasma cells are formed.

#### ZBTB20 deficiency causes overactivity of B cell transcriptional programs

The early functional activity of ZBTB20 led us to explore the cell types in which this transcription factor may act. Immunization with NP-CGG/alum drives the development of recently activated GL7^positive^ CD38^positive^ antigen-specific B blast cells and antigen-specific GL7^positive^ CD38^negative^ GC B cells by 7 days post-immunization **(Figures 1A and S1A**). Together, these cells are precursors to GC B cells, early memory B cells, and plasma cells^48^. NP-specific GC B cell and B blast frequencies were similar in ZBTB20^KO^ and Mb1-Cre+ control mice **(Figures S4A-B).** We therefore isolated B blasts and GC B cells from these mice at 7 days post-immunization and GC B cells at 14 days post-immunization via FACS for single-cell RNA-sequencing (scRNA-seq) to define mechanisms that promote durable antibody responses.

We observed 9 distinct transcriptional clusters across cell types and days post-immunization **(Figures 3A-D)**^56–59^. Using Monocle^60–62^, we observed pseudotime trajectories consistent with known developmental pathways of day 7 B blasts into day 7 GC B and day 14 GC B populations **(Figure S4C)**. Despite the functional requirement for ZBTB20 by day 7 post-immunization, we observed considerable overlap between ZBTB20^KO^ and Mb1-Cre+ control clusters within the day 7 B blasts and GC B cells (**Figure 3D**). Although 91 differentially expressed genes (DEGs) were observed in day 7 ZBTB20^KO^ GCB compared to Mb1-Cre+ controls, only 3 of these genes exhibited at least a 1.5-fold difference, and only two genes showed greater than a 2-fold change **(Supplemental Table 1)**, perhaps explaining the similar clustering patterns (**Figure 3D**). Similarly, bulk RNA-seq revealed that only 12 and 27 genes had at least a 2-fold expression difference between ZBTB20^KO^ and Mb1-Cre+ genotypes in day 7 B blasts and GC B cells, respectively (**Figures S4D-E** and **Supplemental Tables 2-3**). At Day 14, when ZBTB20 is no longer required for durable antibody production **(Figures 2C-D)**, we observed 265 DEGs between ZBTB20^KO^ and Mb1-Cre+ control cells, 22 of which showed at least a 1.5-fold difference in the scRNA-seq data **(Figure 3D** and **Supplemental Table 4)**.

**Figure 3:**
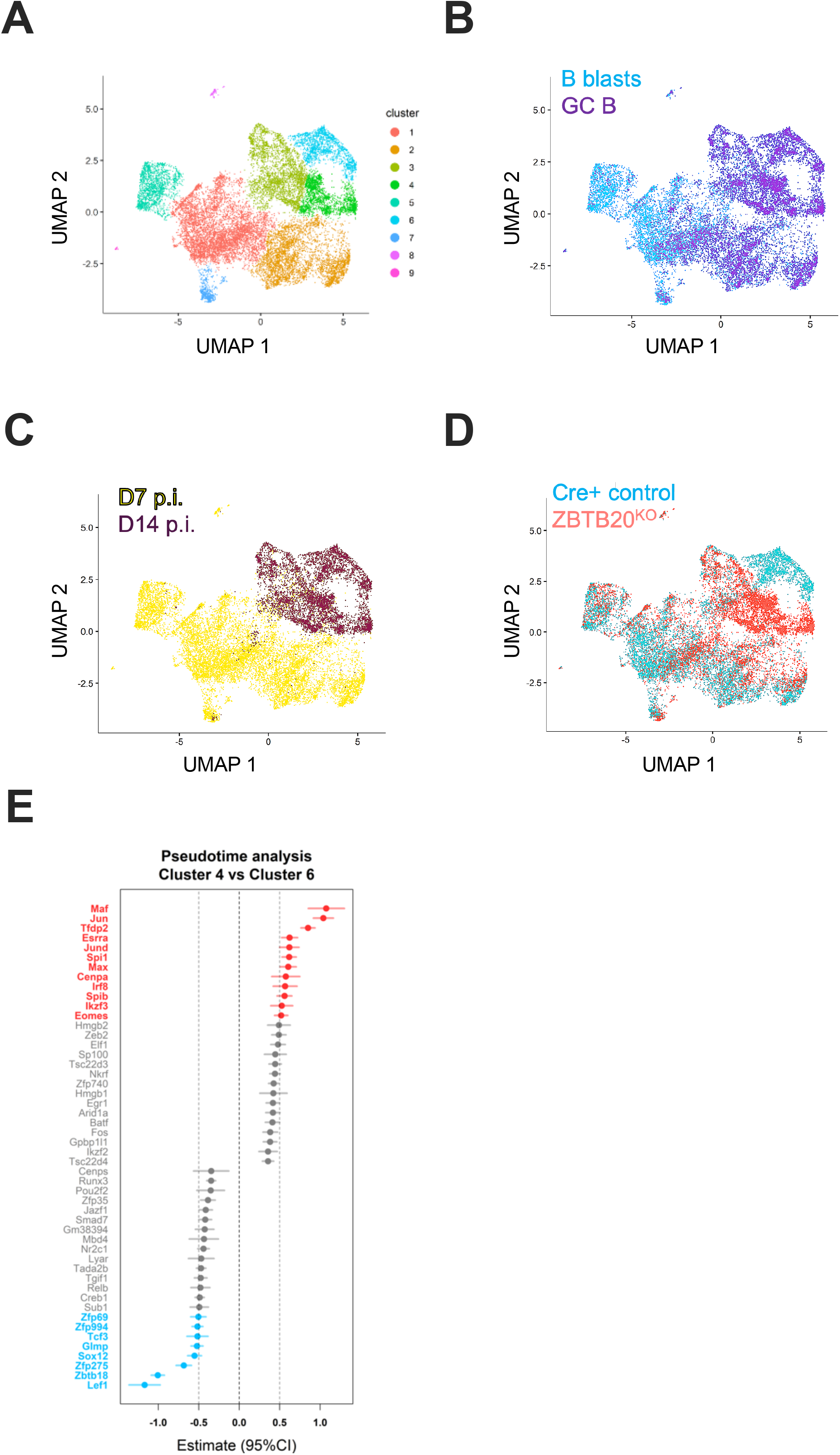
Loss of ZBTB20 in germinal center B cells leads to progressively dysregulated activity of B cell transcriptional programs. (**A-D**) Single-cell transcriptomes of NP-specific B blasts and GC B cells from ZBTB20^KO^ and Cre+ control mice were analyzed using Seurat and Monocle bioinformatic packages. (**A**) Resultant UMAP plot depicting 9 transcriptional clusters across day 7 and day 14 cells is shown. Day 7 results are cumulative of cells from 4 ZBTB20^KO^ and 5 Mb1-Cre+ control mice.Day 14 GC B gene expression data are cumulative of cells from 5 ZBTB20^KO^ and 5 Mb1-Cre+ control mice. (**B**) Clusters in (A) are distinguished based on the identity of the cells sorted for library preparation as B blasts (blue) and GC B cells (purple). (**C**) Clusters were separated by time point post immunization (p.i.), showing cells at day 7 (yellow) and day 14 (dark purple). (**D**) Transcriptional profiles overlaid with ZBTB20^KO^ (red) and Cre+ control (blue) genotypes. (**E**) Gene regulatory network scoring of differential transcription factor activity for cluster 4 of scRNA-seq datasets (enriched for ZBTB20^KO^ day 14 GC B cells) compared to cluster 6 (enriched for Cre+ control day 14 GC B cells) using PISCES.

The magnitude of expression differences in any individual gene was relatively modest, even at Day 14. Therefore, we employed PISCES to gain insight into regulatory programs, rather than individual genes, dysregulated by the absence of ZBTB20.

PISCES constructs gene regulatory networks of transcription factors within single-cell data using the ARACNe and metaVIPER pipelines^63,64^. Comparisons of D14 ZBTB20^KO^-associated cluster 4 and D14 Mb1-Cre+-associated cluster 6 suggested, among others, increased activity of the transcription factor IRF8, the Ets family transcription factors PU.1 (encoded by *Spi1*) and SPIB, and members of the AP-1 family of transcription factors such as Jun and JunD **(Figure 3E)**. Of note, IRF8 can physically interact with Ets and AP-1 family members to promote binding and transcription at Ets-IRF composite elements (EICEs) and AP1-IRF composite elements (AICEs), respectively^65–68^, and antagonizes the plasma cell fate^69^.

#### ZBTB20 deficiency perturbs epigenetic signatures prior to transcriptional effects

Although ZBTB20 is required only within the first week post-immunization, the magnitude and numbers of transcriptional differences between ZBTB20^KO^ and control cells were modest at Day 7 post-immunization. We hypothesized ZBTB20 may impart an epigenetic imprint in early GC B populations that precedes marked impacts on gene expression. To test this hypothesis, we conducted single-cell ATAC-seq (scATAC-seq), which measures accessible chromatin regions across the genome. ZBTB20^KO^ and Mb1-Cre+ control animals were immunized and antigen-specific GC B cells were analyzed by scATAC-seq at 7 days post-immunization and epigenetic analyses were conducted using Signac^70^. Unlike the relatively similar scRNA-seq profiles at this time point, we observed a clear epigenetic divergence between day 7 ZBTB20^KO^ and Mb1-Cre+ control cells **(Figure 4A)**. We detected 485 differentially accessible regions (DAR) between ZBTB20^KO^-associated cluster 1 and Mb1-Cre+-associated cluster 0 **(Figures S5A-B** and **Supplemental Table 5)**. Similar results were observed after integrating and re-clustering the data sets using Harmony to mitigate batch effects **(Figures S5C-E** and **Supplemental Table 6)**^71^.

**Figure 4:**
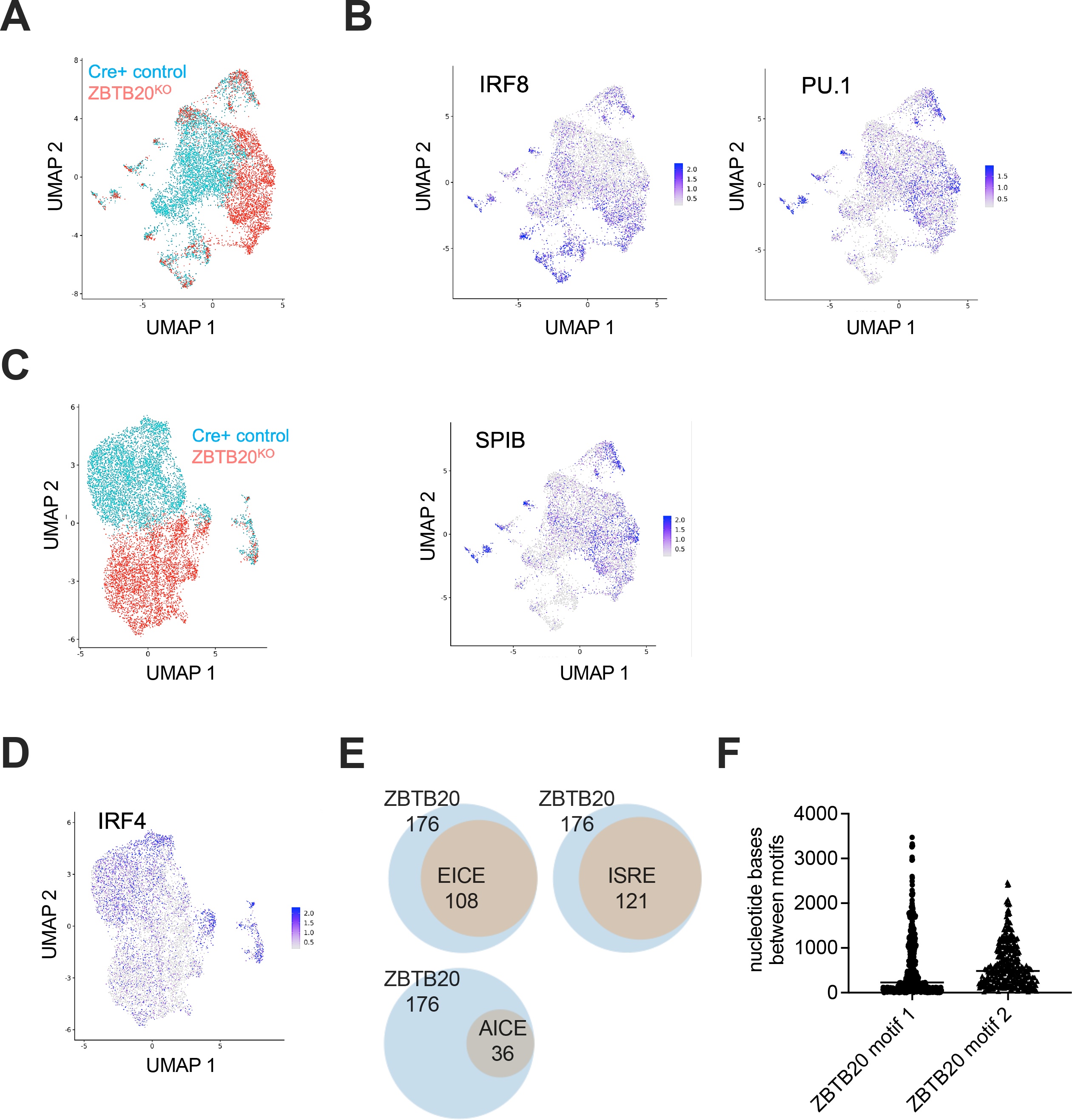
ZBTB20 deficiency perturbs epigenetic programs in early GC B cells. **(A-B)** Single-cell epigenetic profiles of ZBTB20^KO^ and Cre control NP specific GC B cells were assessed at 7 days post-immunization using Signac. Data filtering and clustering methods are detailed in methods sections. Data are cumulative of cells from nine ZBTB20^KO^ and 10 Mb1-Cre+ control mice. (**B**) ChromVAR motif accessibility scoring of transcription factors IRF8 (adj. p-value = 5.12 x 10^-41^), PU1 (adj. p-value = 1.22 x 10^-41^), and SPIB (adj. p-value = 1.01 x 10^-41^) displayed enrichment of consensus binding sites in the ZBTB20^KO^ associated clusters compared to the predominant Cre+ control associated cluster. Full chromVAR scoring for day 7 transcription factors with differential motif availability in primary ZBTB20^KO^ enriched clusters are listed in **Supplemental Table 7**. (**C-D**) Day 14 NP specific GC B cells from ZBTB20^KO^ and Cre+ control mice were examined for chromatin accessibility by scATAC-seq. (C) UMAP plot depicting single-cells from ZBTB20^KO^ (red) and Cre+ controls (blue) are shown. Data are cumulative of cells from 7 ZBTB20^KO^ and 7 Mb1-Cre+ control mice. (D) Differential IRF4 target accessibility (p-value = 5.29 x 10^-61^) were determined by ChromVAR and shown across both genotypes of cells. Full chromVAR scoring for day 14 transcription factors with differential motif availability in primary ZBTB20^KO^ enriched cluster are listed in **Supplemental Table 8.** (**E**) Differentially accessible regions (DARs) from primary ZBTB20^KO^ associated cluster 1 and Cre+ control primary associated cluster 0 were assessed for ZBTB20 consensus binding motifs and composite ISRE, EICE, and AICE binding sites. DARs containing at least one ZBTB20 consensus binding sites are indicated in blue circles in the Venn diagrams, and co-occurrences with other binding motifs are displayed in orange. (**F**) The distance between ZBTB20 binding motifs and consensus ISRE binding sites within DARs are shown. Bars indicate the median distance. scATAC-seq libraries were generated from sorted antigen-specific GC B cells.

To identify candidate transcription factors driving epigenetic dysregulation in the absence of ZBTB20, we employed chromVAR, which predicts differential transcription factor activity in scATAC-seq data using known DNA binding motifs in promoters and enhancers^72,73^. The binding sites of 405 transcription factors were enriched in differentially accessible regions (DARs) between the dominant ZBTB20^KO^ clusters and the major Mb1-Cre+ control-associated cluster **(Supplemental Table 7)**. Of particular interest, IRF8, PU.1, and SPIB activities were predicted by chromVAR to be highly enriched in ZBTB20^KO^ cells **(Figure 4B)**. These three transcription factors were also independently predicted by PISCES to have elevated activity at day 14 post-immunization from the scRNA-seq data above (**Figure 3E**). We next tested whether the epigenetic dysregulation in ZBTB20^KO^ cells was sustained as the GC progressed. We conducted scATAC-seq analysis of ZBTB20^KO^ and Mb1-Cre+ control GC B cells at day 14 and day 21 post-immunization. We observed continued ZBTB20-dependent epigenetic dysregulation across these times **(Figures 4C** and **S5F).** ChromVAR scoring in day 14 GC B cells indicated a subtle but statistically significant decrease in the activity of the plasma cell lineage-promoting and survival transcription factor IRF4 in ZBTB20^KO^ relative to control cells **(Figure 4D** and **Supplemental Table 8)**. Cumulatively, these results demonstrate ZBTB20 imprints an early epigenetic signature that may impact the longevity of plasma cells formed much later.

To gain further insight into how ZBTB20 deficiency leads to elevated Ets factor and IRF8 activity, we first quantified protein levels of PU.1 and IRF8 in ZBTB20^KO^ and Mb1-Cre+ control NP+ GC B cells at 7 days post-immunization. No statistically significant differences were observed between genotypes (**Figures S5G-H**). We next quantified potential ZBTB20 binding sites within DARs of the primary ZBTB20^KO^ associated and Mb1-Cre+ control associated scATAC-seq clusters from antigen-specific GC B cells 7 days post-immunization. Because the ZBTB20 antibody failed in ChIP-seq and Cut&Run assays, we instead queried DARs, which had a median size of 1155bp, for the two 15 bp consensus ZBTB20 binding sites identified from previous overexpression ChIP-Seq experiments^74^. Potential ZBTB20 binding sites within DARs were identified using the FIMO program from the MEME-suite^75^. Of 485 DARs between the Mb1-Cre+ control-associated cluster 0 and ZBTB20^KO^-associated cluster 1 (**Figure S5A**), 176 had at least one predicted ZBTB20 binding site (**Figure 4E** and **Supplemental Table 9**). None of these were located near the *Irf8* or *Spi1* loci. Of these 176 DARs with consensus ZBTB20-binding site sequences, 121 also contained at least one ISRE sequence (**Figure 4E** and **Supplemental Table 10**), which are low-affinity sites bound by IRF dimers^67^. Similarly, we observed that 108 DARs contained both ZBTB20 consensus binding sites and EICE sequences, which are bound by IRF-Ets heterodimers (**Figure 4E** and **Supplemental Table 10**)^65,76^. Less overlap (36 DARs) was observed between ZBTB20 consensus binding sites and AICE sequences, which are bound by heteromeric complexes of IRF and AP-1 factors (**Figure 4E** and **Supplemental Table 10**)^65,76^. Within the DARs, ISRE sites were located at median distances of 228bp and 487bp from consensus ZBTB20 binding motifs 1 and 2, respectively (**Figure 4F**), arguing against direct physical competition for the same binding sites. These data suggest that ZBTB20 may repress IRF8 and Ets targets by rendering chromatin inaccessible.

The scATAC-seq data suggested impaired activity in ZBTB20^KO^ GC B cells of IRF4, a transcription factor required for both plasma cell formation and longevity^67,77,78^. To determine if ZBTB20 deficiency restricts the formation and migration of GC-derived plasma cells to the bone marrow, we employed Bromodeoxyuridine (BrdU) pulse-chase experiments. We immunized ZBTB20^KO^ and Mb1-Cre+ control mice with NP-CGG/alum and provided them with BrdU-containing drinking water from 3.5 weeks to 4 weeks post-immunization. We identified BrdU+ antigen-specific plasma cells in the bone marrow, which represent recently formed cells derived from proliferating precursors **(Figure 5A)**. No statistically significant differences were observed in BrdU+ antigen-specific bone marrow plasma cell frequencies and total numbers between ZBTB20^KO^ and control mice **(Figures 5B-D)**. However, we observed slightly reduced protein levels of IRF4 in BrdU+ plasma cells of ZBTB20^KO^ mice compared to Mb1-Cre+ controls **(Figure 5E)**. These data suggest that while plasma cells are formed at normal rates, they may express insufficient levels of IRF4.

**Figure 5:**
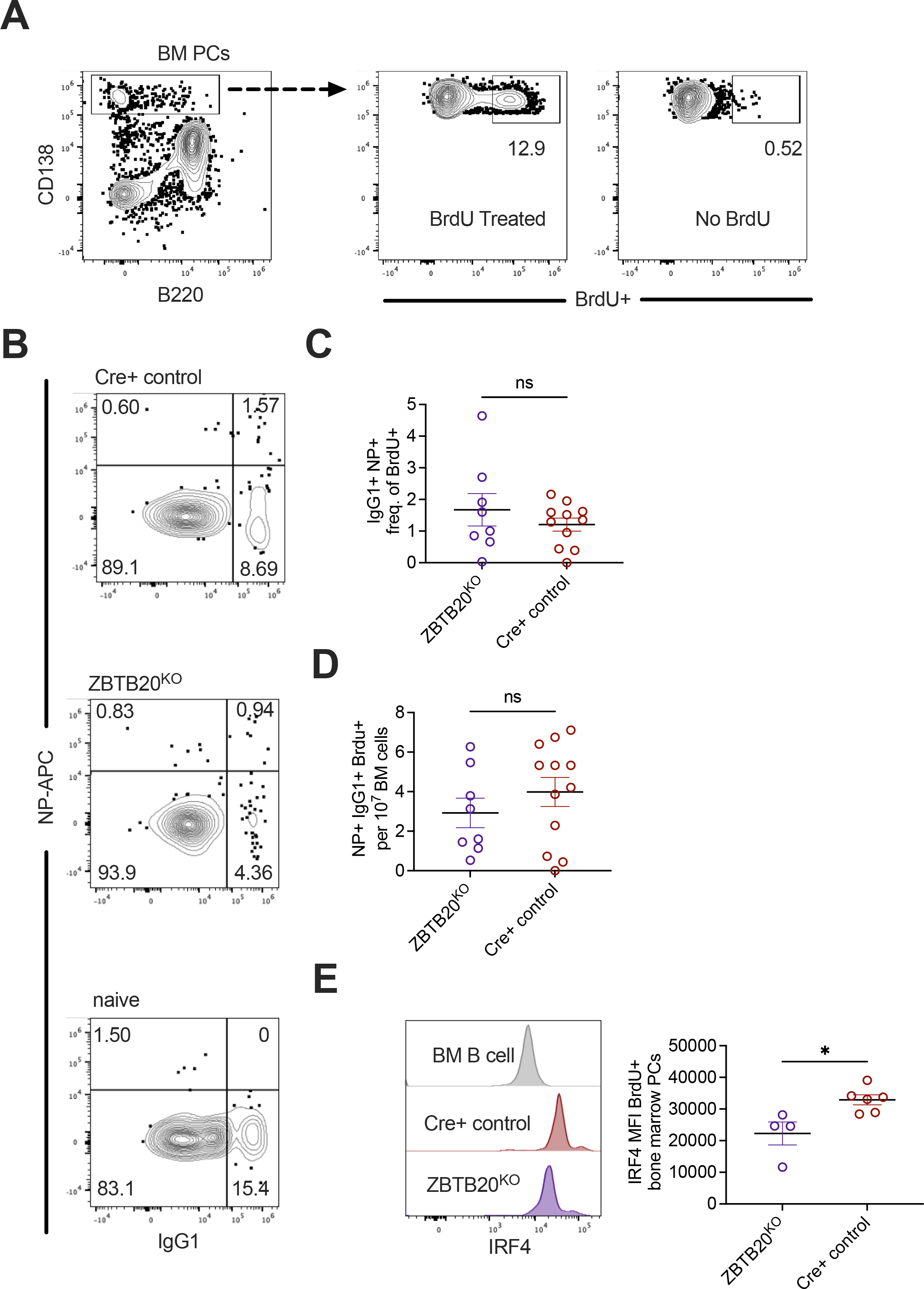
ZBTB20 deficient plasma cells are formed at normal rates but have diminished IRF4 expression. (**A**) Representative flow cytometry of bone marrow plasma cells labeled by BrdU treated drinking water from 3.5 weeks to 4 weeks following NP-CGG/alum immunization. (**B**) Frequencies of double positive NP+ IgG1+ bone marrow plasma cells within the BrdU+ fraction were quantified using the gating strategy in (A). (**C**) Frequencies of NP+ IgG1+ cells within BrdU+ plasma cells across genotypes were quantified for ZBTB20^KO^ (purple) and Cre+ control (red) mice. Each dot represents an individual mouse and data shown are representative of two individual experiments. No statistical significance was observed by an unpaired two-tailed t-test. (**D**) Total numbers of NP+ IgG1+ BrdU+ cells were quantified per 10 million bone marrow cells for groups in (C). Each dot represents an individual mouse. Data are representative of two individual experiments. Mean values ± SEM are shown. No statistical significance was observed by unpaired student’s two-tailed t-test. (**E**) Intracellular IRF4 protein was quantified by flow cytometry in bone marrow BrdU+ plasma cells of ZBTB20^KO^ (purple) and Cre+ control (red) mice. Bone marrow B cells (grey) were used as a control. Each dot represents an individual mouse. Data are representative of two individual experiments. Mean values ± SEM are shown. *p< 0.05 by unpaired student’s two-tailed t-test.

#### Suppression of IRF8 and Ets factor activity promotes plasma cell longevity

Previously, we observed that the oil-in-water Sigma adjuvant, which is similar to Ribi, partially rescued durable antibody production in ZBTB20 gene-trapped fetal liver chimeras^36^. This rescue was recapitulated in ZBTB20^KO^ mice immunized with NP-CGG/Sigma adjuvant **(Figures 6A-B)**. To determine whether this rescue might occur through the suppression of IRF8, Ets, and/or AP–1-dependent epigenetic programs, we isolated antigen-specific GC B cells from ZBTB20^KO^ and Mb1-Cre+ control mice immunized with NP-CGG/Sigma adjuvant and analyzed them by scATAC-seq. Although scATAC-seq analysis still demonstrated an epigenetic divergence between ZBTB20^KO^ and Mb1-Cre+ control mice immunized with NP-CGG/Sigma adjuvant **(Figure 6C)**, IRF8, PU.1, and SPIB activity as inferred by ChromVAR was not statistically different between genotypes **(Figures 6D, S6A,** and **Supplemental Table 11).** Yet when examining ZBTB20^KO^ GC B cells, we found that PU.1 and SPIB activity was enriched when alum was used in the immunization compared with Sigma adjuvant **(Figures 6E-F, S6B-C,** and **Supplemental Table 12)**. These data provide circumstantial evidence that repression of Ets transcription factor activity is important for durable antibody production.

**Figure 6:**
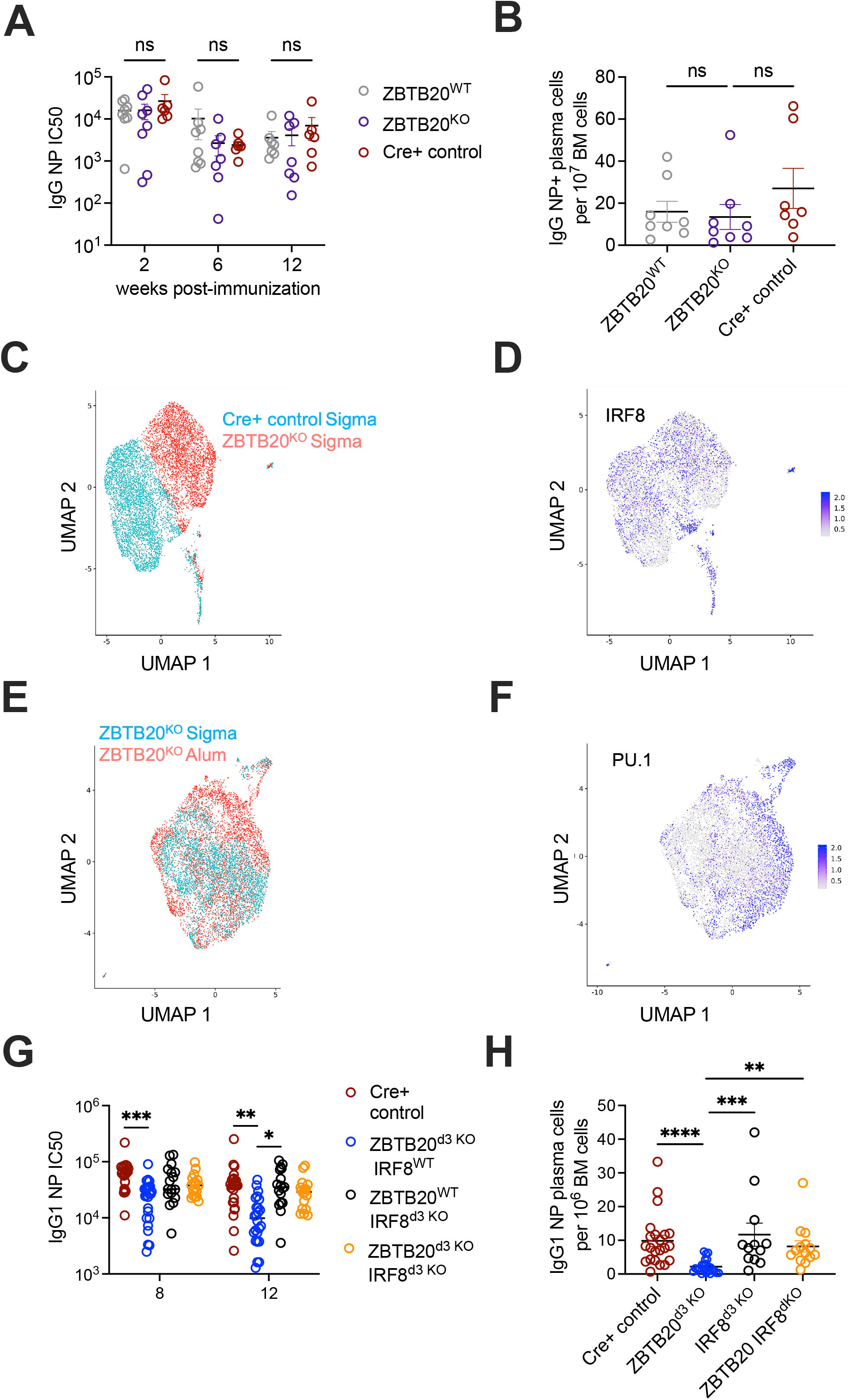
Ets and IRF8 programs antagonize plasma cell longevity. **(A-B)** ZBTB20^KO^ mice were immunized with NP-CGG/Sigma adjuvant alongside littermate and age matched Cre+ controls. (A) IgG anti-NP serum antibodies were quantified via ELISA in ZBTB20^KO^ (purple), ZBTB20^WT^ (grey), and age-matched Cre+ controls (red). The IC50 values for individual mice were derived from serial dilution curves at each time point indicated. Each dot represents an individual mouse. Data are cumulative of two independent experiments. Mean values ± SEM are shown. Two-way ANOVA with mixed-effects model and Tukey multiple comparisons testing of IC50 values showed no significant differences across genotypes across the indicated time points. (B) IgG NP-specific bone marrow plasma cells were quantified via ELISPOT at 12 weeks post-immunization in ZBTB20^KO^, ZBTB20^WT^, and age-matched Cre+ controls. Each dot represents an individual mouse. Data are cumulative of two independent experiments. Mean values ± SEM are shown. No statistical significance was determined by Kruskal-Wallis test with Dunn’s multiple comparisons. (**C**) ZBTB20^KO^ and Cre+ control mice were immunized with NP-CGG/Sigma adjuvant and antigen-specific GC B cells were isolated 14 days post-immunization for scATAC-seq. UMAP clusters are separated by genotype into ZBTB20^KO^ (red) and Cre+ controls (blue). Day 14 NP-CGG/alum samples are identical to those in Figure 4. For additional day 14 NP-CGG/Sigma adjuvant libraries, data are cumulative of cells from 9 ZBTB20^KO^ and 9 Mb1-Cre+ control mice. (**D**) Cells with accessible loci enriched for IRF8 binding sites are indicated among cells in (C). ChromVAR scoring for IRF8 showed no statistically significant differences between genotypes. (**E**) Single-cell epigenetic analysis of day 14 GC B cells from ZBTB20^KO^ mice immunized with NP-CGG/Alum (red) or NP-CGG/Sigma adjuvant (blue) were plotted for epigenetic profiles and chromVAR transcription factor binding motif availability. (**F**) Cells with accessible loci enriched for PU.1 binding sites across both genotypes are shown for cells in (E). PU.1 binding sites in DARS are enriched in ZBTB20^KO^ alum associated cells (adj. p-value = 5.43 x 10^-129^). (**G**-**H**) Zbtb20^fl/fl^ Irf8^fl/fl^ hCD20 CreERT2+/- double knockout mice were immunized with NP-CGG/Alum alongside single knockout controls for each transcription factor and Cre+ controls. Tamoxifen was gavaged 3 days post-immunization in all animals. **(G)** NP+ IgG1+ antibody titers were quantified for Cre+ control (red), hCD20 CreERT2+/- *Zbtb20*^fl/fl^ (blue), hCD20 CreERT2+/- *Irf8*^fl/fl^ (black), and double knockout mice (orange). Each dot represents an individual mouse. Data are cumulative of four independent experiments. Mean values ± SEM are shown. *= p-value <0.05, ** = p-value < 0.01, *** = p-value < 0.001, by two-way ANOVA with mixed-effects model and Tukey multiple comparison testing. (**H**) IgG1 NP bone marrow plasma cells were quantified for mice in (G) via ELISPOTs at 12 weeks post-immunization. Each dot represents an individual mouse. Data are cumulative of four independent experiments. Mean values ± SEM are shown. *= p-value <0.05, ** = p-value < 0.01, *** = p-value < 0.001, by Kruskal-Wallis test with Dunn’s multiple comparisons.

To more directly test the functional consequences of these elevated transcriptional programs, we crossed hCD20 CreERT2+ mice with *Zbtb20^fl/fl^* and/or *Irf8^fl/fl^* mice, as IRF8 interacts with PU.1. Mice were immunized with NP-CGG/alum and oral tamoxifen was administered 3 days post-immunization. As expected, NP-specific antibody levels were lower in ZBTB20^d^^3^ ^KO^ mice relative to hCD20 CreERT2+ controls at 8 and 12 weeks post-immunization **(Figure 6G).** Deficiency of IRF8 alone led to titers similar to those of hCD20 CreERT2+ controls and elevated relative to ZBTB20^d^^3^ ^KO^ mice at 12 weeks post-immunization **(Figure 6G)**. At weeks 8 and 12, deficiency of both ZBTB20 and IRF8 led to NP-specific antibody titers that were not statistically different from those of the hCD20 CreERT2+ and IRF8^KO^ groups, suggesting at least a partial rescue of ZBTB20-deficiency in durable antibody production **(Figure 6G)**. At 12 weeks post-immunization, NP-specific plasma cells as measured by ELISPOT were elevated in ZBTB20^KO^ IRF8^KO^ mice relative to ZBTB20^KO^ animals, but similar to hCD20 CreERT2+ and IRF8^KO^ controls **(Figure 6H)**, again suggesting a functional rescue. Together, these data suggest that adjuvant- and IRF8-responsive epigenetic programs are established early in the B cell response and imprint plasma cell longevity.

## DISCUSSION

In our prior work using *Zbtb20* gene-trapped or germline knockout chimera mice, the defects in durable antibody production seemed specific to long-lived plasma cells^36,44^. We observed no ZBTB20-dependent changes in GC B cell numbers or organization, affinity maturation, memory B cell numbers, short-lived plasma cell numbers, or the rate of plasma cell formation. For these reasons, we assumed that ZBTB20 functionally acted directly in long-lived plasma cells, perhaps by subtly influencing anti-apoptotic protein levels, to promote survival and durable immunity. Yet in light of our current work, this interpretation needs to be revised. Our studies instead revealed perhaps a more interesting fundamental principle, which is that subtle epigenetic programs are established very early during B cell activation to imprint the survival of plasma cells that are formed much later. These programs are largely hidden from view since the rest of the B cell response is seemingly unaffected by their dysregulation and precede major impacts on transcription. In this sense, the key conclusions of our study have less to do with ZBTB20 itself than the epigenetic and transcriptional programs that its absence revealed.

Still, some key questions remain regarding how ZBTB20 functions. Given that ZBTB20 likely acts as a repressor^47,79,80^, the *Irf8* or *Ets* loci themselves would be obvious candidates for direct binding. However, we did not observe consensus ZBTB20 binding sites as differentially accessible in the promoters or enhancer regions for these genes, nor did we observed altered protein levels of these factors. Instead, we found that ZBTB20 deficiency leads to altered accessibility at many regions that contain both predicted ZBTB20 binding sites and ISRE and/or EICE sequences. Thus, ZBTB20-mediated repression of a subset of IRF8 and Ets targets may occur by rendering chromatin in these areas inaccessible. This subtle dysregulation is maintained and potentially amplified into later germinal center B cells, which normally preferentially produce long-lived plasma cells^17,19^. We have not been able to obtain enough downstream knockout antigen-specific plasma cells to profile and fully determine the consequences of the upstream germinal center transcriptional dysregulation. Yet consistent with prior studies of IRF8 antagonism in B cells and other lineages^66,69^, we did observe slightly reduced IRF4 protein levels in ZBTB20-deficient plasma cells. Given the known role of IRF4 in maintaining long-lived plasma cells^81,82^, this may help explain how ZBTB20 indirectly impacts survival.

Notably, the innate signals triggered by the immunization impacted these early epigenetic programs, as an oil-in-water adjuvant similar to Ribi was more effective at suppressing Ets-dependent programs in ZBTB20-deficient B cells than was alum. Perhaps as a result, ZBTB20-deficient antibody responses were rescued when this oil-in-water adjuvant was employed. In our study, we employed an oil-in-water adjuvant that also contains TLR ligands. Yet in prior studies using West Nile virus veterinary vaccines, we observed that oil-in-water adjuvants that lack TLR ligands were also able to rescue ZBTB20 deficiency^36^, whereas in the current work, addition of the TLR ligand CpG was insufficient to rescue ZBTB20-deficient antibody responses. These data suggest that the oil-in-water component of the adjuvants, rather than TLR ligands, was the important variable in promoting survival of ZBTB20-deficient plasma cells. The sensors, target cell types, and mechanisms of action of oil-in-water adjuvants are not well understood^83^, but they likely act on myeloid cells rather than on B cells directly. Thus, the B cell epigenetic programs are likely influenced by adjuvants indirectly.

Prior studies in wild-type mice have shown that the duration of NP-specific antibody production is similar in alum vs. Ribi-adjuvanted responses^35^. In preliminary experiments, we have confirmed that the durability of NP-specific responses in wild-type backgrounds is similar between alum and Sigma adjuvants, but we have also observed adjuvant-dependent differences in the durability of antibody production when protein antigens such as SARS-CoV-2 Spike are used as the immunogen, consistent with recent non-human primate studies^34^. Thus, there seems to be a complex interplay between the antigen and adjuvant in instructing durable responses that might potentially converge on programs such as those driven by Ets and IRF8 transcription factors. Engineering vaccines towards such key early epigenetic signatures may enable rapid improvements in durability without necessarily waiting for years to see how each change fared.

## ACKNOWLEDGEMENTS

This work was supported by NIH grants R01AI099108 (D.B.), R01AI129945 (D.B.), R21AI176305-01A1 (A.B.), R35GM137896 (D.A.C.), and P30CA023074 (for the Flow Cytometry Shared Resource). This work was also supported by a Gates Foundation grant INV-051356 (D.B.). We are also grateful to the University of Arizona Genomics Core for use of the Bioanalyzer. T.J.R. was supported by the ARCS Foundation and L.J.D. was supported by a fellowship from the BIO5 Institute. We thank Rachel Wong and Jennifer Uhrlaub for mentorship in laboratory immunological assays, and Hao Zhang and Holly Welfley for help with single-cell epigenetic analyses.

## METHODS

### Experimental Resources and Models

#### Lead Contact

Further information and reagents or resources requests should be directed to and will be fulfilled by the lead contact, Dr. Deepta Bhattacharya.

#### Animal models

Zbtb20^fl/fl^ mice were previously described^50^. Live Irf8^fl/fl^ mice were a gift from Dr. Harinder Singh at the University of Pittsburgh, and are available from the Jackson Laboratory (JAX) as strain #:014175 RRID:IMSR_JAX:014175^84^. Mb1Cre+ mice are available from JAX as Strain #:020505 RRID:IMSR_JAX:020505^49^. J-Chain IRES CreERT2 mice are available from JAX as Strain #:035764 RRID:IMSR_JAX:035764^52^. hCD20 TamCre+ mice were a gift from Dr. Mark Shlomchik’s laboratory^33^.

#### Data and Code Availability

Sequencing data are available on the NCBI SRA database, Accession Code PRJNA1292783.

##### Methods Details

#### Mice

All experimental mice were housed and bred in pathogen-free facilities at the University of Arizona Animal Health Science Center. All animal procedures used were approved by the Animal Care and Use Committees at the University of Arizona (protocol 17-266). Sex and age matched, 7-16-week-old female and male mice were equally assigned to experimental groups. Cre+ mice were bred with either *Zbtb20*^fl/fl^ or *Irf8*^fl/fl^ mice and then intercrossed until mice were homozygous for the floxed alleles. Breeding pairs were maintained such that litters were a mix of Cre^+/-^ and Cre^-/-^ mice. Separately, Cre+ control animals were maintained and used as age-matched controls. For timed deletions, mice were orally gavaged once with 4 mg of tamoxifen dissolved in sunflower seed oil and then maintained on chow containing 400mg/kg of tamoxifen (Envigo) for two weeks.

#### Flow Cytometry and Sorting

Following euthanasia, spleen and/or bone marrow was prepared for single-cell suspension. All steps were performed using 1x PBS with 5% equaFetal Bovine Serum (Atlas Biological) as sample buffer unless otherwise noted. Red blood cells were lysed using ammonium chloride potassium lysis buffer. Cells were further separated from debris by density gradient centrifugation by overlaying 2mL sample with equal volume of Histopaque-1119 (Millipore Sigma) or Lympholyte (Cedarlane Labs). Lymphocytes were then stained with antibody panels on ice and placed away from light. Relevant antibodies for this study are listed in the key resource table with the indicated fluorophores, clone, and vendor. NP-APC was generated in the lab as described previously^85^. To assess intracellular levels of transcription factors, cells were fixed, permeabilized, and intracellularly stained following the True-Nuclear Transcription Factor Buffer Set (Biolegend 424401). Samples for single-cell sequencing analysis were enriched with anti-PE microbeads (Miltenyi Biotech 130-048-801).

Enriched samples were resuspended in RPMI 1640 with 10% FBS and sorted into RPMI with 25% FBS to assist viability of sorted cells. All flow cytometric analysis was conducted on machines provided and maintained by the University of Arizona Flow Cytometry Core. Fluorescent activated cell sorting was performed using the FACS Aria III (BD Biosciences). Flow analysis for protein quantification was performed on either the LSRII (BD Biosciences) or the Aurora (Cytek Biosciences). Data processing and quantification was performed utilizing FlowJo v.10 and GraphPad Prism v.9 and v.10.

#### ELISAs

For NP serum ELISAs of mice models, 96 well flat bottom plates (Corning) were coated with NP-BSA at 5 µg/mL in 0.1M sodium bicarbonate buffer pH 9.6 overnight at 4 degrees C. Following overnight incubation, all subsequent steps were performed at room temperature for one hour and washed a minimum of three times with PBS + 0.05% Tween (PBS-T) between each step. Serum and antibodies were diluted in PBS + 2% adult bovine serum (ABS). Coated plates were blocked with 1x PBS + 2% ABS for one hour. Serum samples were serial diluted 10-fold beginning with an initial concentration of 1:100 for NP-CGG/alum immunized mice. For NP-CGG/Sigma Adjuvant immunized mice, samples were diluted 4-fold beginning with a 1:50 dilution.

Blocked plates were washed and overlayed with serial serum dilution series. Serum antibodies were probed with anti-mouse IgG1 biotin (BD Pharmigen 553441) at 50ng/mL for NP-CGG/alum immunizations and Jackson ImmunoResearch IgG biotin at 100ng/mL for NP-CGG/Sigma immunizations. Secondary antibodies were probed with Streptavidin-HRP (BD Biosciences 554066) at a dilution factor of 1:1000. Following PBS-T and PBS wash, antibody bound signal was detected using 100uL Alfa Asear 3,3’,5,5’-Tetramethylbenzidine solution high sensitivity. Plates were developed for 30 seconds before quenching with 50uL 2N H2SO4. Quantification of Optical Density was done using a colorimeter at the 450nm wavelength. IC50 values of serial dilution curves were determined in GraphPad Prism using regression of OD 450nm signal with a 0.05 technical background serving as the baseline.

WNV E D-III ELISAs were performed as above except for the coating antigen and the serial dilution was 4-fold beginning with a 1:50 dilution of serum.

#### ELISPOTs

Multiscreen HTS 96 well filter plates (Millipore Sigma) were incubated overnight at 4 degrees with 10 µg/mL of NP-BSA (LGC Biosearch Technologies in 1x PBS. Bone marrow lymphocytes were harvested as described in the Flow Cytometry and Sorting section. ELISPOT plates were washed with three times with 1x PBS and twice with PBS-T and blocked for 1 hour at room temperature with RPMI containing 10% FBS. Following blocking, plates were washed again and bone marrow samples were plated in triplicate at 1 million cells per well in RPMI + FBS for NP-CGG/alum immunized animals. For NP-Sigma and WNV-E DIII immunizations bone marrow samples, CD138+ cells were enriched from bone marrow samples using magnetic enrichments after staining with anti-mouse CD138 PE (Biolegend) followed by anti-PE microbeads (Miltenyi Biotec). Five percent of the bone marrow sample was preserved for flow cytometric analysis to determine pre-enrichment plasma cell frequencies. Ten percent of the enriched sample was preserved to determine enriched cell counts and assessed by flow cytometric analysis to determine post-enrichment plasma cell frequencies. The remaining enriched sample was plated for ELISPOT.

ELISPOT plates with plated cells were incubated overnight at 37^0^C. Incubated plates were washed vigorously with 1x PBS by multichannel pipetting. NP-CGG/alum and WNV E-DIII/alum immunized samples were probed with anti-mouse IgG1 biotin (BD Pharmigen) at 50ng/mL and NP-CGG/Sigma samples were probed with anti-mouse IgG biotin (Jackson ImmunoResearch) at a concentration of 2.5 µg/mL. Plates were then washed again and incubated with SA-HRP (BD Pharmingen) at a dilution factor of 1:5000 for one hour away from light. Following washing, ELISPOT plates were developed with 50uL of TrueBlue Peroxidase Substrate (Kierkegaard and Perry Laboratories) for 5 minutes at room temperature. Plates were then rinsed in distilled water and dried overnight before spots were enumerated using CTL ImmunoSpot S6 software.

#### WNV Focus Reduction Neutralization Tests

Serum samples from WNV E DIII-CpG/alum immunized mice were heat inactivated by one hour incubation at 65 degrees. Vero CC81 cells (ATCC) were plated in 96 well plates at a density of 2 million cells per plate and incubated overnight. WNV-NY99 (gift of M. Diamond, Washington University) was incubated at a concentration of 100 pfu per well with serially diluted serum from immunized mice for two hours. Following incubation, cells were treated with 1.5% methylcellulose (Millipore Sigma) in DMEM with 5% FBS and left at 37^0^C for 48 hours. Plates were then flooded in 1% paraformaldehyde (PFA) solution and left at room temperature for 1 hour to fix. Following a wash with BD Perm Wash Buffer (BD Biosciences) infected cells were probed for using the pan flavivirus antibody hE16 (BEI Resources) at a concentration of 2.5µg/mL in BD Perm Wash Buffer and incubated at room temp for 1 hour. An anti-human IgG HRP secondary (Jackson ImmunoResearch) was diluted to a concentration of 1:2000 in BD Perm Wash Buffer and incubated for one hour. Plates were then washed and incubated with TrueBlue Peroxidase Substrate for 10 minutes and rinsed with distilled water. Foci were counted using the CTL ImmunoSpot S6 software.

#### Mouse Immunizations

Mice were immunized intraperitoneally with 100µg of NP-CGG conjugated in 5% aluminum potassium sulfate (alum) adjuvant or sigma adjuvant system (Millipore Sigma) as described previously^36^. WNV E-DIII immunizations were performed intraperitoneally with 25µg protein mixed with 20µg of the mouse TLR9 ligand ODN-1826 and conjugated in alum adjuvant^52,86^.

#### BrdU Treatment and Quantification

At 3.5 weeks post immunization, mice were given drinking water containing dissolved BrdU (Sigma Aldrich) at a concentration of 0.8mg/mL. At 4 weeks post-immunization, bone marrow samples were stained with an anti-mouse CD138 PE antibody (Biolegend) and enriched with anti-PE magnetic microbeads (Miltenyi Biotec). Following enrichment on LS columns (Miltenyi Biotec), samples were processed using the FITC BrdU Flow Kit (BD Biosciences) per the manufacturer’s instructions. Cells were stained with NP-APC and anti-mouse IgG1 (Biolegend) while staining for BrdU to identify antigen-specific plasma cells. Cells were enumerated as a ratio of total NP+ IgG1+ CD138+ bone marrow plasma cells per 10M total bone marrow cells harvested, or as the frequency of NP+ IgG1+ cells of the BrdU+ plasma cell fraction.

#### Analysis of single-cell epigenetic libraries

Sorted antigen-specific GC B cells were processed for Nuclei Isolation using the EZ Lysis Nuclei Isolation Protocol (Millipore Sigma) and buffer exchanged with Diluted Nuclei Buffer (10x Genomics) prior to transposition. For each timepoint, two biological replicate pools of antigen-specific cells from immunized ZBTB20^KO^ and Mb1-Cre+ control mice were generated. For day 7 scATAC-seq analysis, nine ZBTB20^KO^ and ten Mb1-Cre+ control mice were used. For day 14 NP-CGG/alum libraries, seven ZBTB20^KO^ and seven Mb1-Cre+ control mice were used. For day 14 NP-CGG/Sigma libraries, nine ZBTB20^KO^ and nine Mb1-Cre+ control mice were used. For day 21, seven ZBTB20^KO^ and eight Mb1-Cre+ control mice were used. The 10x Single-cell Multiomics protocol was followed library construction.

Sequencing was done at Novogene in Sacramento, California. Fastq files were analyzed by Cell Ranger ATAC 2.0.1 to assess cellular recoveries and initial sample quality metrics. Peak files, meta data tables, fragments and fragments indices were utilized for Signac analysis where further QC and filtering metrics were applied. Baseline filtering metrics were as follows:

Peak region fragments < 250000 &

Pct reads in peaks > 50 &

Nucleosome signal < 1 &

TSS.enrichment > 3

Appropriate dimensionality and clustering metrics were determined empirically using Elbow plots and Depth Correlation plots. GC B cells were further defined by accessibility of the lineage defining transcription factor *Bcl6*. DAR analysis was conducted using FindMarkers of the peak assay with Seurat objects, min.pct thresholds were set at 0.05 and latent.vars = ‘nCount_peaks’ to filter low present reads. chromVAR assessment was followed using the publicly available tutorial https://stuartlab.org/signac/articles/motif_vignette.html using the fold-change calculation parameters to determine average differences in z-scores across compared clusters.

ChromVAR^72^ scoring was done by utilizing known transcription factor binding motifs from the publicly available cis-BP and JASPAR databases^74^ and by scoring accessibility of the given motifs across the entire dataset. ChromVAR z-scores assign the contribution of peaks from individual cells and queried clusters to score the enrichment potential transcription factor binding sites.

Two consensus ZBTB20 binding motifs were extracted from published work using murine CD8 T cells^74^. Differentially accessible regions (DARs) were identified from day 7 post-immunization antigen-specific GC B cells of ZBTB20^KO^ and Cre+ control dominated clusters. Specifically, clusters 0 and 1 were used from **Figure S5A**. DARs were scanned using FIMO^65^ from the MEME Suite for the presence of two ZBTB20 binding motifs (motif 1 = GGAGGCAGAGGCAGG, motif 2 = TGCTGGGATTAAG). A maximum adjusted p-value of 0.01 was applied to select high confidence potential binding sites from ZBTB20 consensus motifs. Potential ZBTB20 binding DARs were then assessed for the presence of ISRE, EICE, and AICE motifs using FIMO and overlap using Microsoft Access^65,67,76^. Venn diagrams were generated using https://www.meta-chart.com/venn#/data.

#### Analysis of transcriptomic libraries

Single cell gene expression libraries were generated from sorted antigen-specific B blasts or GC B cells. Day 7 B blasts and day 7 GC B gene expression libraries were generated from pooled antigen-specific cells sorted from four ZBTB20^KO^ and five Mb1-Cre+ control mice of mixed sexes using the 10X Genomics protocol for 3’ Gene Expression. Day 14 GC B gene expression libraries were generated from pooled antigen-specific cells sorted from five male ZBTB20^KO^ and five male Mb1-Cre+ control mice using the 10X Genomics protocol for 5’ Gene Expression and VDJ isolation. Single-cell library QC was performed using the Agilent High Sensitivity DNA Kit (Agilent Technologies) and assessed for size distribution using an Agilent bioanalyzer 2100 housed in the University of Arizona Genomics Core.

Samples were sequenced at Novogene in Sacramento, California. Sequenced reads were assessed for gene expression using the 10X Cell Ranger 6.0.1 pipeline and mapped back to the mm10 reference transcriptome. Output files were imported into the R statistical environment (R version 4.1.2) for quality control assessment. Briefly, barcodes without a corresponding gene expression profile were identified and removed with the emptyDrops function from the DropletUtils package, and genes were filtered to retain those expressed in at least 0.1% of cells. Cells were excluded from the analysis if they met any of the following conditions: had less than 500 total genes identified, had less than 1000 transcripts expressed, had greater than 50% ribosomal gene content, or had a mitochondrial gene content greater than an adaptive threshold (median + 4X the median absolute deviation). Genes located on the X or Y chromosomes were excluded from both Day 7 and Day 14 analyses since cells from both sexes were pooled in the Day 7 samples. The data was converted to a Seurat object and the standard Seurat pipeline (Seurat Version 3.2.3) was applied (e.g, the NormalizeData, FindVariableFeatures, ScaleData, RunPCA, FindNeighbors, FindClusters, RunUMAP, and CellCycleScoring functions). Finally, SingleR^75^ employing the ‘Immgen’ and ‘mouseRNA’ databases, was applied for unbiased cell type annotation to confirm B cell identity of the single-cell profiles generated. Differentially expressed genes were detected with seurat’s FindMarkers function employing the MAST framework^87^. Genes were considered differentially expressed if they exhibited a Bonferroni-adjusted p-value < 0.05 and an average Log2 fold change of +/- 0.25.

For PISCES analysis, 500 metacells were created for each cluster and ARACNe was run with a list of known mouse transcription factors. Regulons are generated through mutual information indexing of transcription factor expression and their inferred target genes paired across the single-cell matrix. Thresholds for statistical significance are set based on the number of instances the pair is observed in the input. Indirect interactions are removed by applying a data processing inequality tolerance filter^50^. Consensus networks for each regulon were generated by estimating statistical significance of the number of times a transcription factor-target pair is detected across the dataset based on a Poisson distribution. Only networks with a Bonferroni-adjusted p-value of < 0.05 are retained^51^. Following regulon constructions, VIPER was employed to score transcription factor regulon activity across the single-cell dataset^76^. Developmental trajectories for scRNA-seq were constructed using Monocle3^62^. Day 7 B blasts were selected as the starting node for Monocle trajectories.

For Bulk RNA sequencing, antigen-specific GC B cells and B blasts were isolated 7 days post-immunization via FACS and RNA was collected using the NuceloSpin RNA Isolation Kit (Machery-Nagel). RNA was sent to Novogene (Sacremento, CA) for low-input RNA-seq library preparation and following library preparation sequencing was performed. Fastq files were aligned to the mm10 reference genome, transcript reads were quantified, and differential expression was calculated across genotypes and cell types on the Galaxy server (usegalaxy.org) using Salmon^88^, DESeq2^89^, and HISAT2^90^.

#### Quantification of Statistical Significance

Statistical calculations for functional experiments in mouse models were performed in GraphPad Prism 10. The specific test employed for each experiment are indicated in figure legends. Single-cell sequencing analyses employed the R (version 4.1.2) programming language for statistical analysis.

##### Key resources table

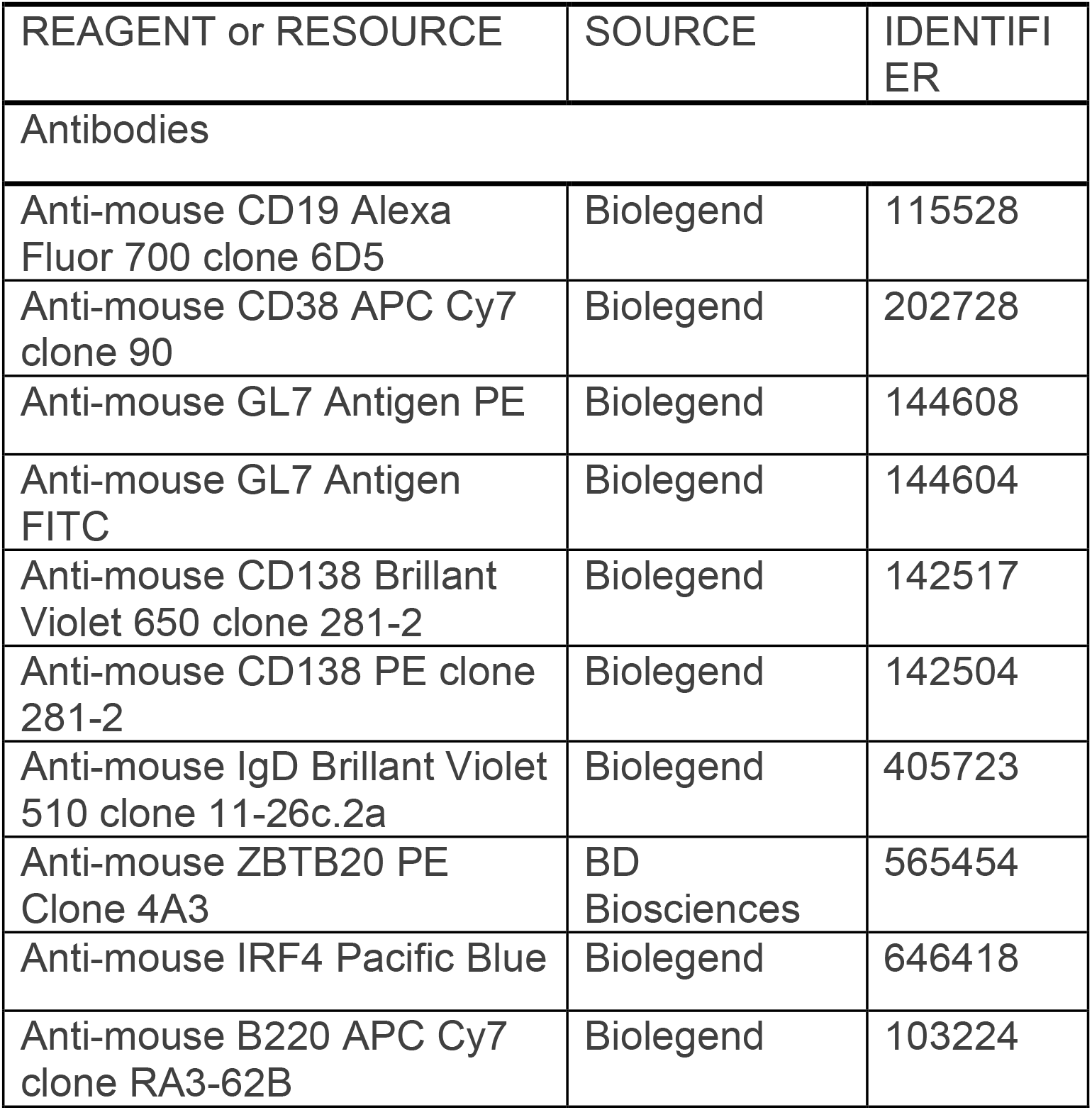
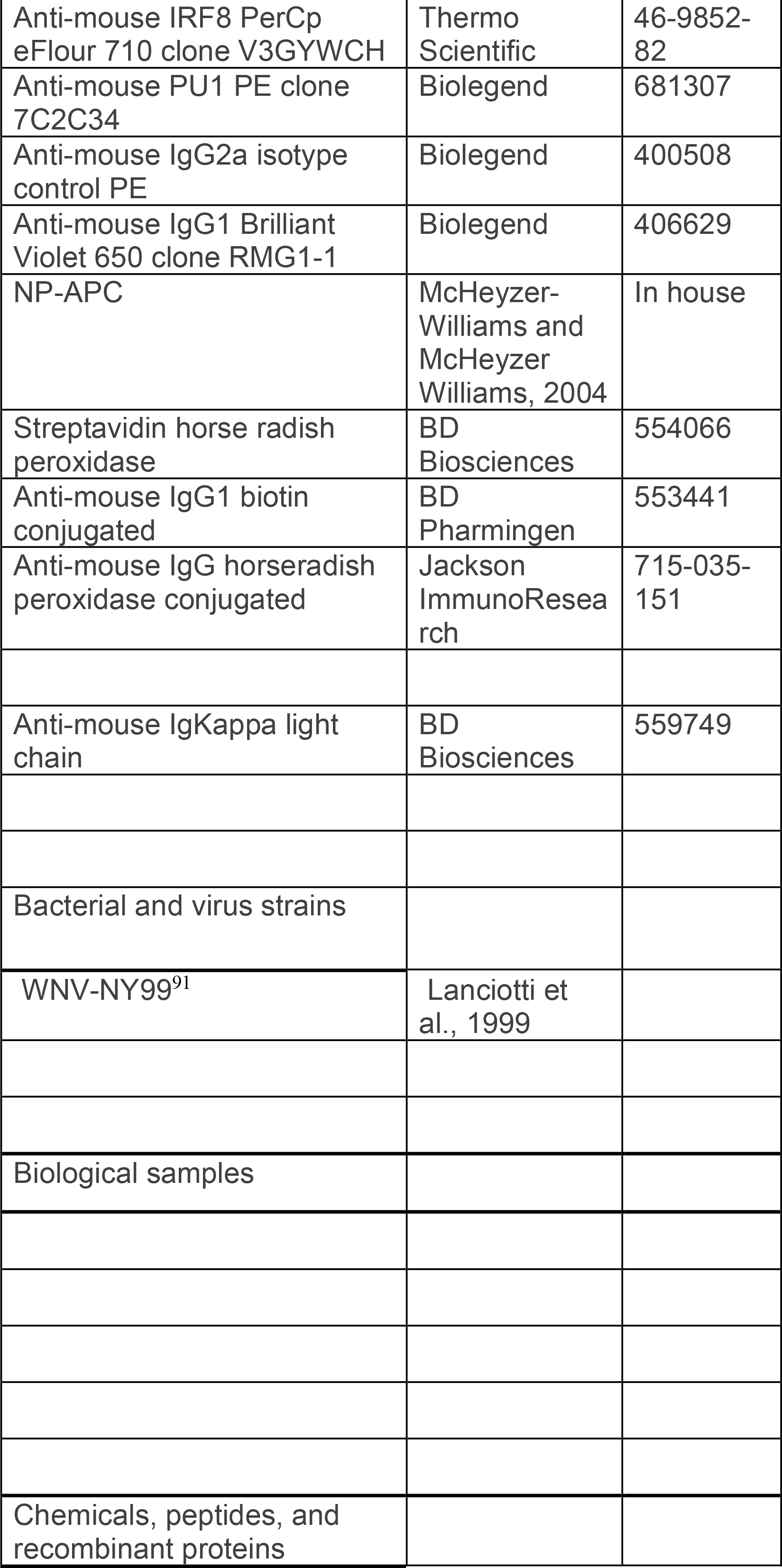
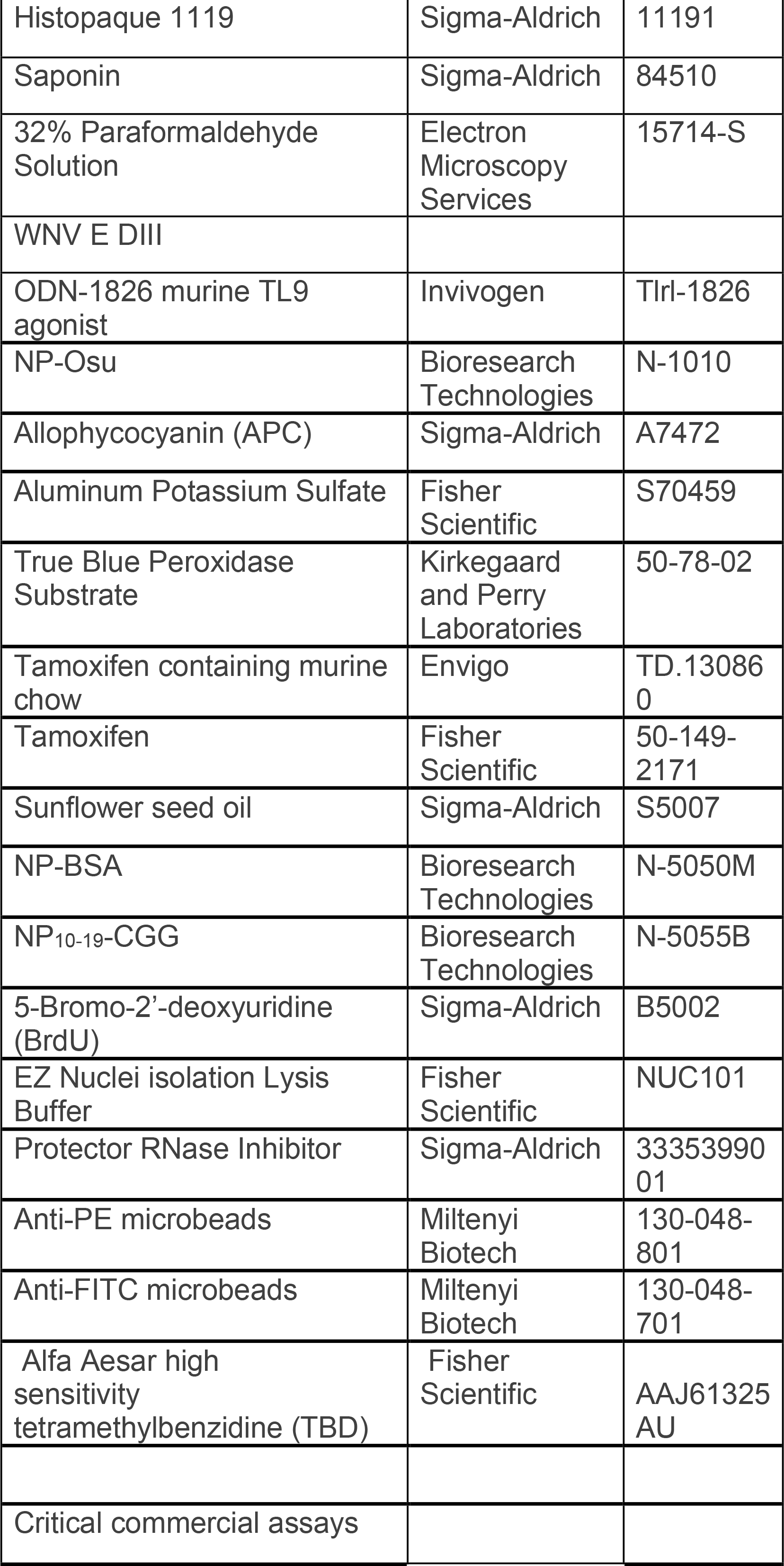
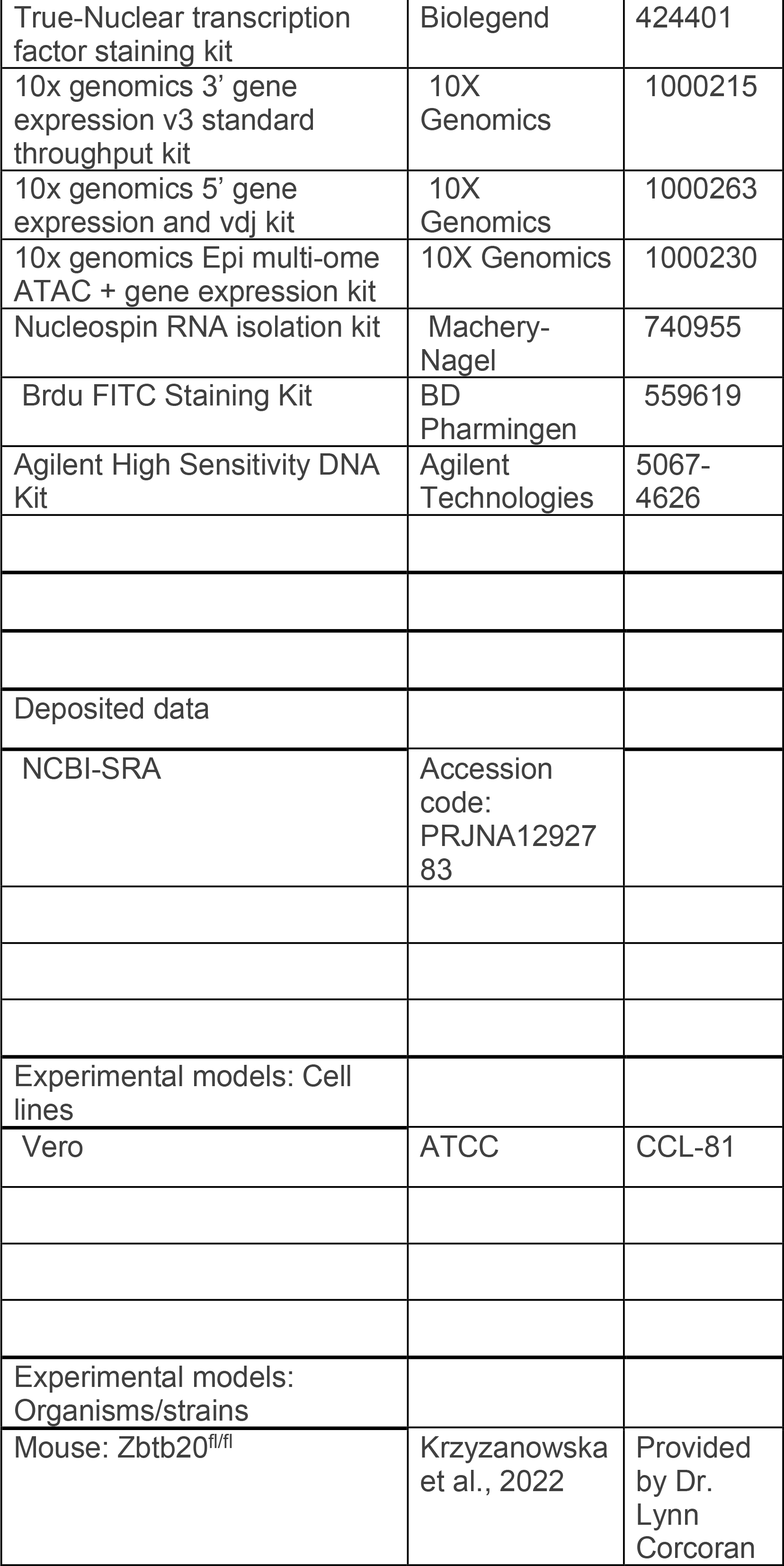
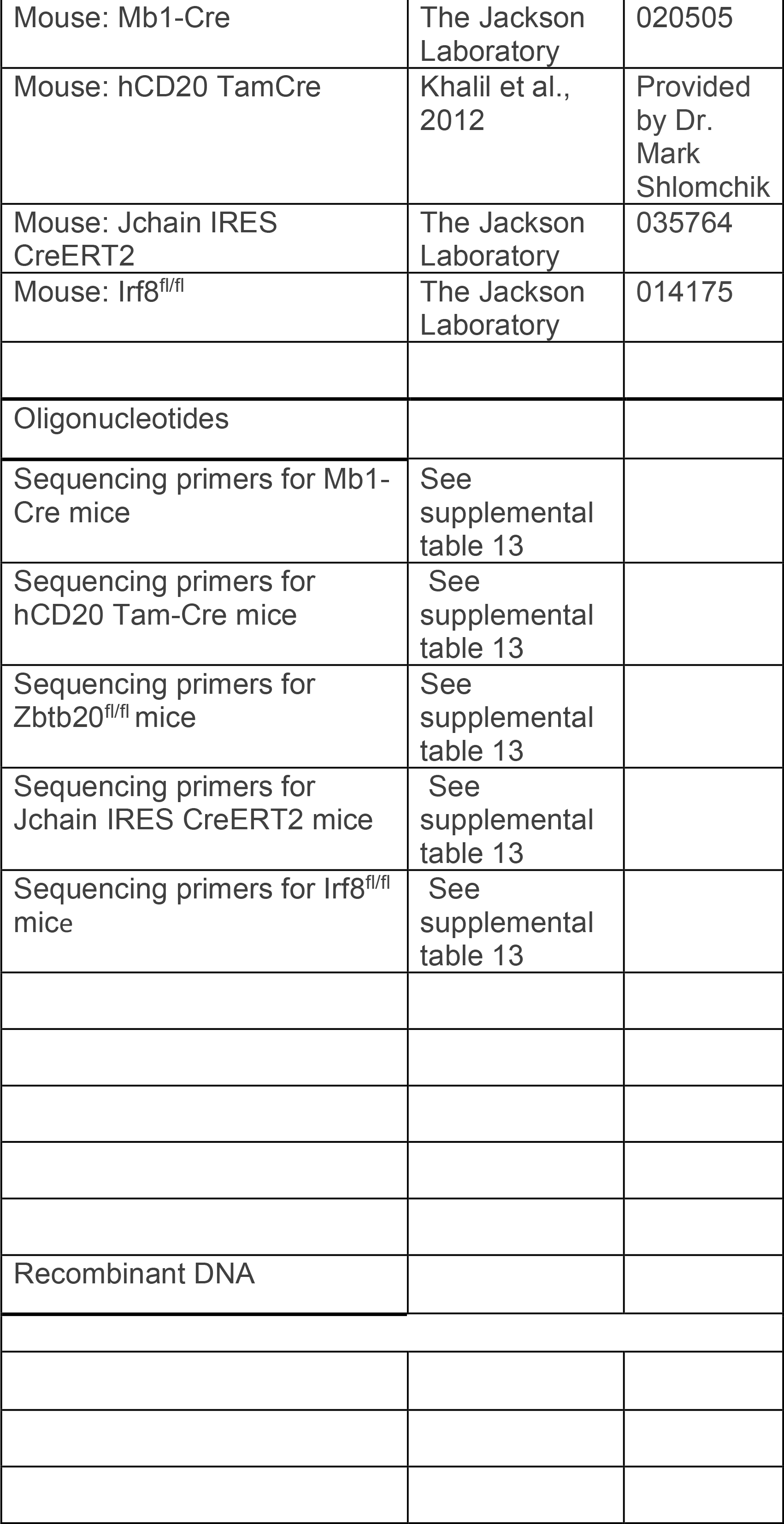
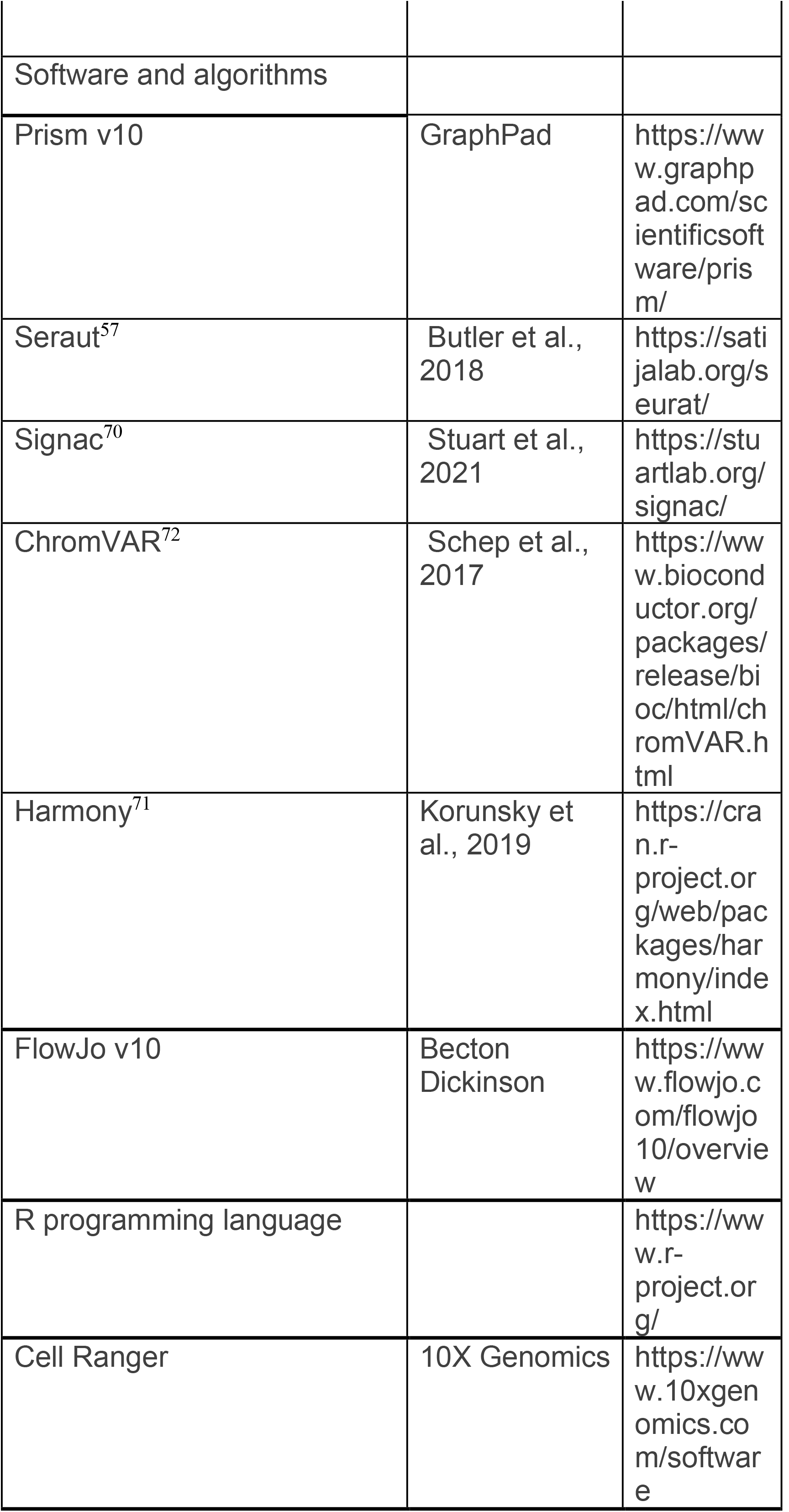
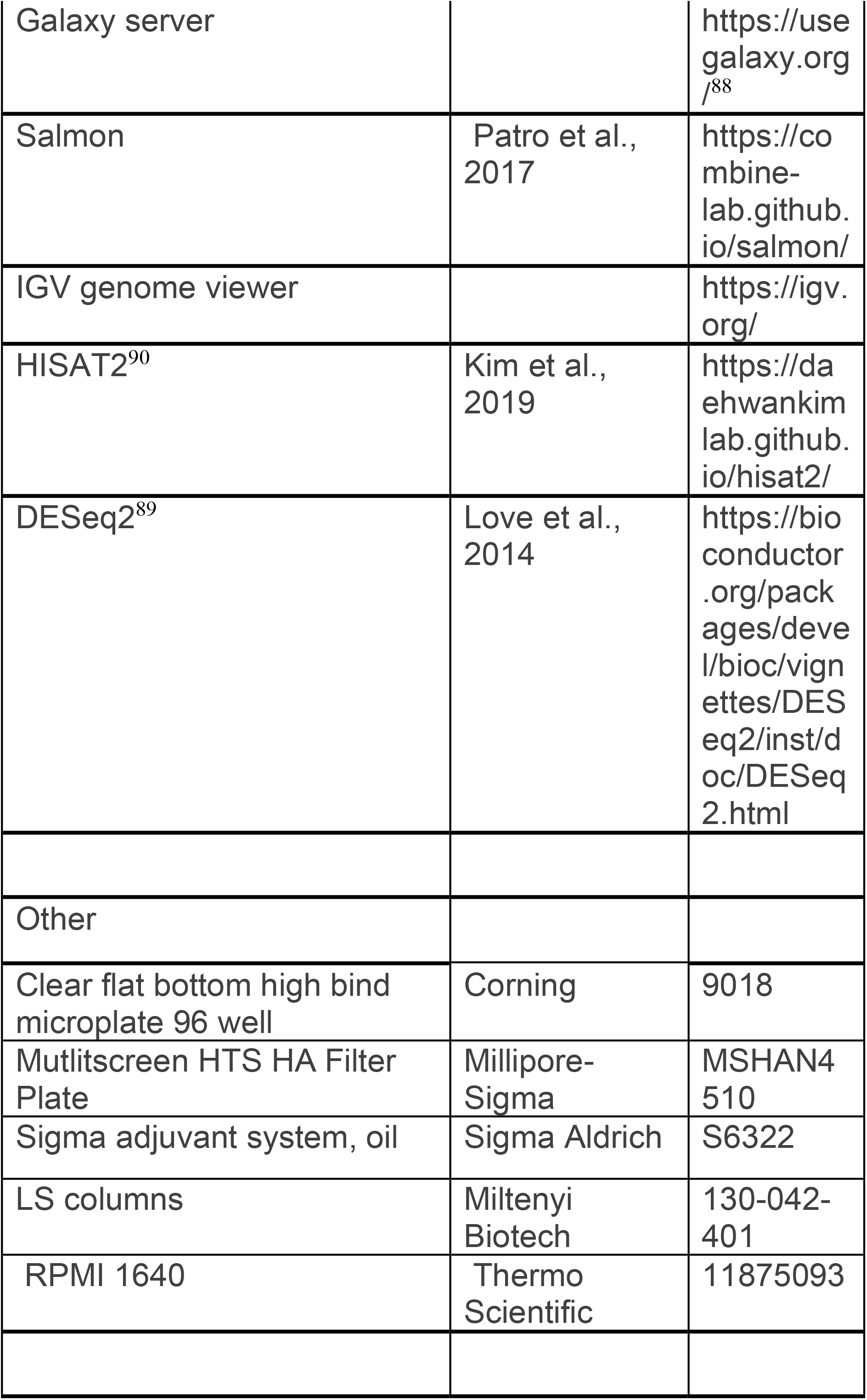

---

**Figure S1 related to Figure 1:**
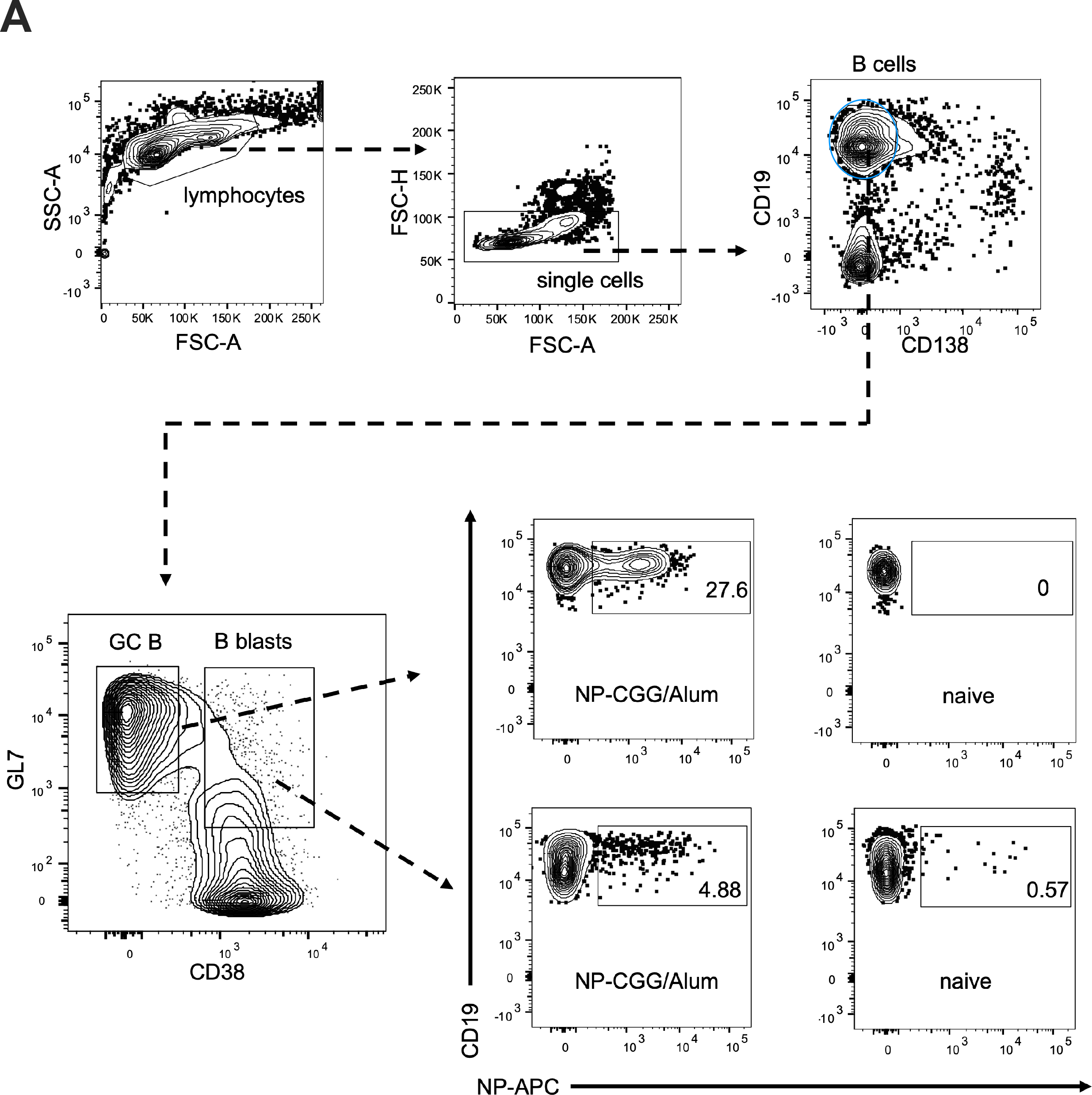
Gating Strategy for antigen-specific GC B and B blasts. (**A**) Representative flow cytometric plots from 7 days post-immunization in ZBTB20^WT^ mice immunized intraperitoneally with NP-CGG/alum.

**Figure S2 related to Figure 1:**
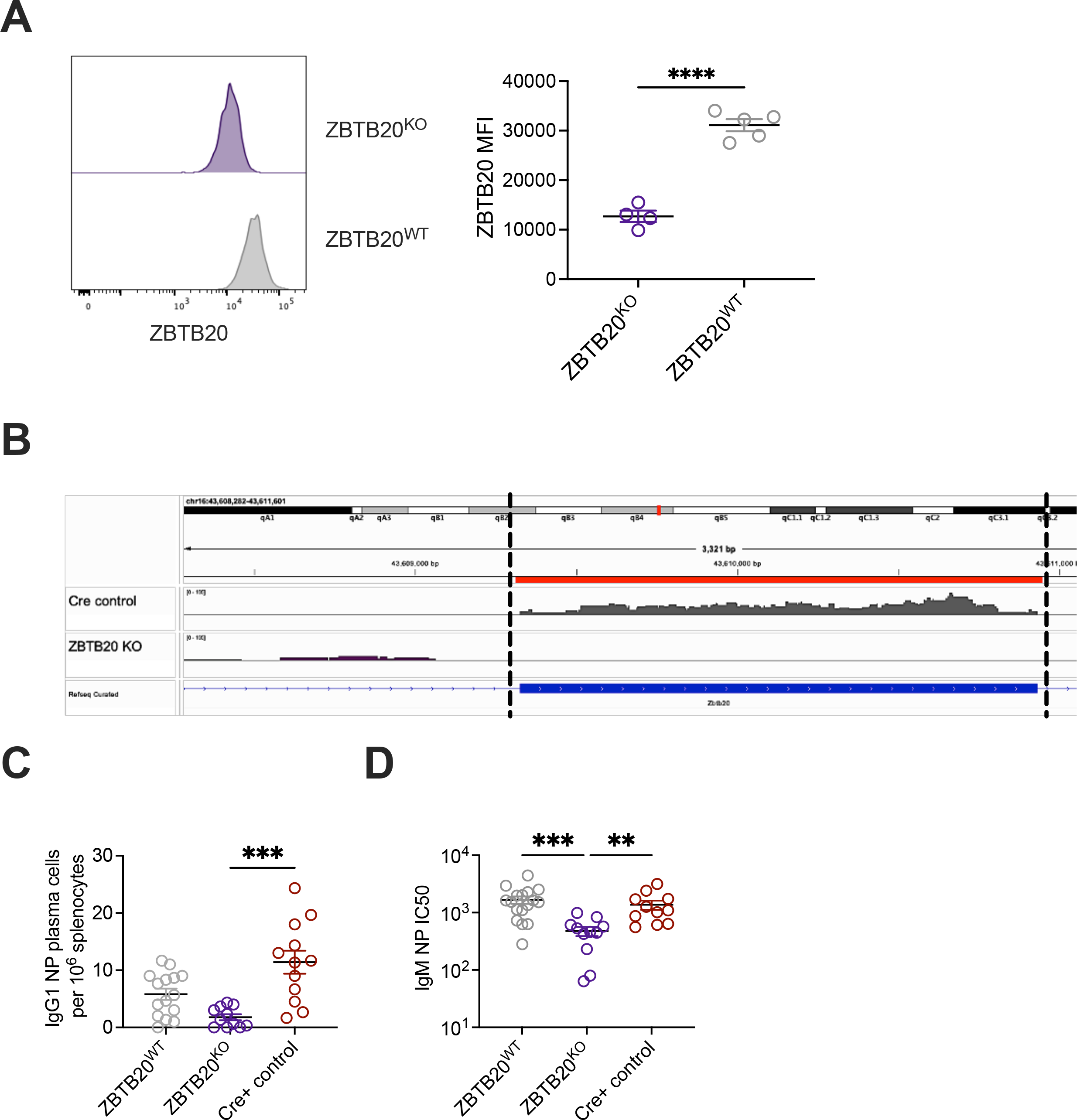
Mb1-Cre depletes *Zbtb20* transcript and protein and restricts IgM+ plasma cell responses. Related to Figure 1. (**A**) ZBTB20^KO^ (purple) and littermate wild type controls (grey) were immunized with NP-CGG/alum and ZBTB20 protein was quantified two weeks post-immunization in antigen-specific GC B cells by intracellular flow cytometric staining. Representative histograms (left) from each animal group are depicted and ZBTB20 MFI in NP specific GC B cells are shown (right). Each dot represents an individual mouse. Data are representative of three individual experiments. Mean values ± SEM are shown. *= p-value <0.05, ** = p-value < 0.01, *** = p-value < 0.001, **** = p-value < 0.0001, by unpaired student’s two-tailed T-test. (**B**) Transcript reads were visualized by RNA-seq conducted on NP-specific GC B cells from ZBTB20^KO^ (purple) and ZBTB20^WT^ (grey) mice 7 days post NP-CGG/alum immunization. Transcript reads were processed using the Salmon and HiSAT2 bioinformatics tools and bam files were aligned for visualization to the mm10 reference genome *Zbtb20* exon 14 (transcript: Zbtb20-204 ENSMUST00000114694.8) using Integrated Genome Viewer. (**C**) Splenic NP specific plasma cells were assessed 12 weeks post-vaccination in ZBTB20^KO^ (purple), ZBTB20^WT^ (red), and Cre+ control (grey) mice by ELISPOT. Each dot represents an individual mouse. Data are cumulative from three individual experiments. Mean values ± SEM are shown *= p-value <0.05, ** = p-value < 0.01, *** = p-value < 0.001, by Kruskal Wallis with Dunn’s multiple comparisons testing. (**D**) NP-specific IgM antibody titers were assessed 14 days post-immunization by serum ELISAs. IC50 values were extrapolated from serial dilution curves of individual mice. Each dot represents an individual mouse. Data are cumulative from three individual experiments. Mean values ± SEM are shown. *= p-value <0.05, ** = p-value < 0.01, *** = p-value < 0.001, by Kruskal Wallis with Dunn’s multiple comparisons testing of IC50 values for each condition.

**Figure S3 related to Figure 2:**
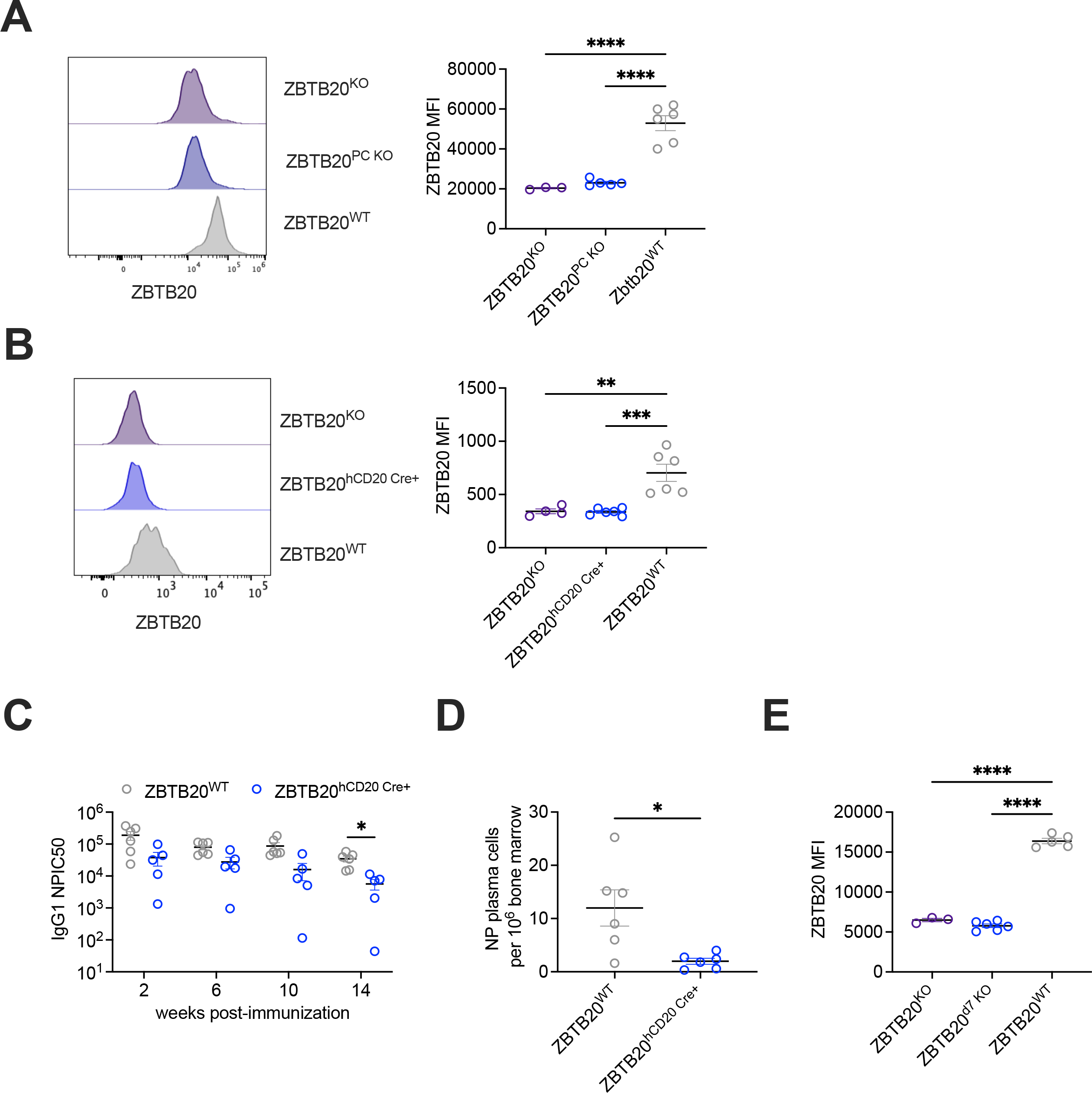
ZBTB20 protein is depleted by tamoxifen inducible Cre systems. (**A**) ZBTB20 protein levels were quantified by flow cytometry in bone marrow plasma cells of ZBTB20^PC KO^ mice (blue) and ZBTB20^WT^ control mice (grey) two weeks following tamoxifen treatment. Plasma cells from ZBTB20^KO^ mice (purple) were utilized as biological controls of ZBTB20 depletion. Representative histograms from each group are depicted (left) and quantified (right). Each dot represents an individual mouse. Data are representative of three individual experiments. Mean values ± SEM are shown. Analysis was conducted using the Cytek Aurora cytometer. *= p-value <0.05, ** = p-value < 0.01, *** = p-value < 0.001, **** = p-value < 0.0001, by one-way ANOVA with Tukey multiple comparisons test for each condition. (**B**) NP specific GC B cells from ZBTB20^hCD^^20^ ^Cre+^ mice were assessed for ZBTB20 protein levels by intracellular flow cytometry four days following tamoxifen gavage. Representative expression histograms from each animal group are depicted (left) and quantified (right). Each dot represents an individual mouse. Data are representative of three individual experiments. Mean values ± SEM are shown. *= p-value <0.05, ** = p-value < 0.01, *** = p-value < 0.001, by 1-way ANOVA with Tukey multiple comparisons test for each condition. Analysis was conducted using the BD LSR II cytometer. (**C**) ZBTB20^hCD^^20^ (blue) and littermate ZBTB20^WT^ control (grey) mice were administered tamoxifen two weeks prior to NP-CGG/alum immunization. NP specific serum ELISAs were conducted longitudinally (C). Each dot represents an individual mouse. Data are representative of two individual experiments. Mean values ± SEM are shown. * = p-value <0.05 by two-way ANOVA with mixed-effects model for IC50 values of each genotype across indicated time points. (**D**) NP specific bone marrow plasma cells were assessed via ELISPOT from mice in (C). Each dot represents an individual mouse. Data are representative of two individual experiments. Mean values ± SEM are shown. * = p-value < 0.05 by unpaired Welch’s two-tailed T-test. (**E**) ZBTB20^hCD^^20^ ^Cre+^ mice were administered tamoxifen 7 days post-immunization (ZBTB20^d^^7^ ^KO^) and deletion was confirmed in antigen-specific GC B cells on day 10 by intracellular flow cytometry. ZBTB20 MFI for NP-specific GC B cells are shown. Analysis was conducted using the Cytek Aurora cytometer. Each dot represents an individual mouse. Data are representative of three individual experiments. Mean values ± SEM are shown. *= p-value <0.05, ** = p-value < 0.01, *** = p-value < 0.001, **** = p-value < 0.0001, by one-way ANOVA with Tukey multiple comparisons test for each condition.

**Figure S4 related to Figure 3:**
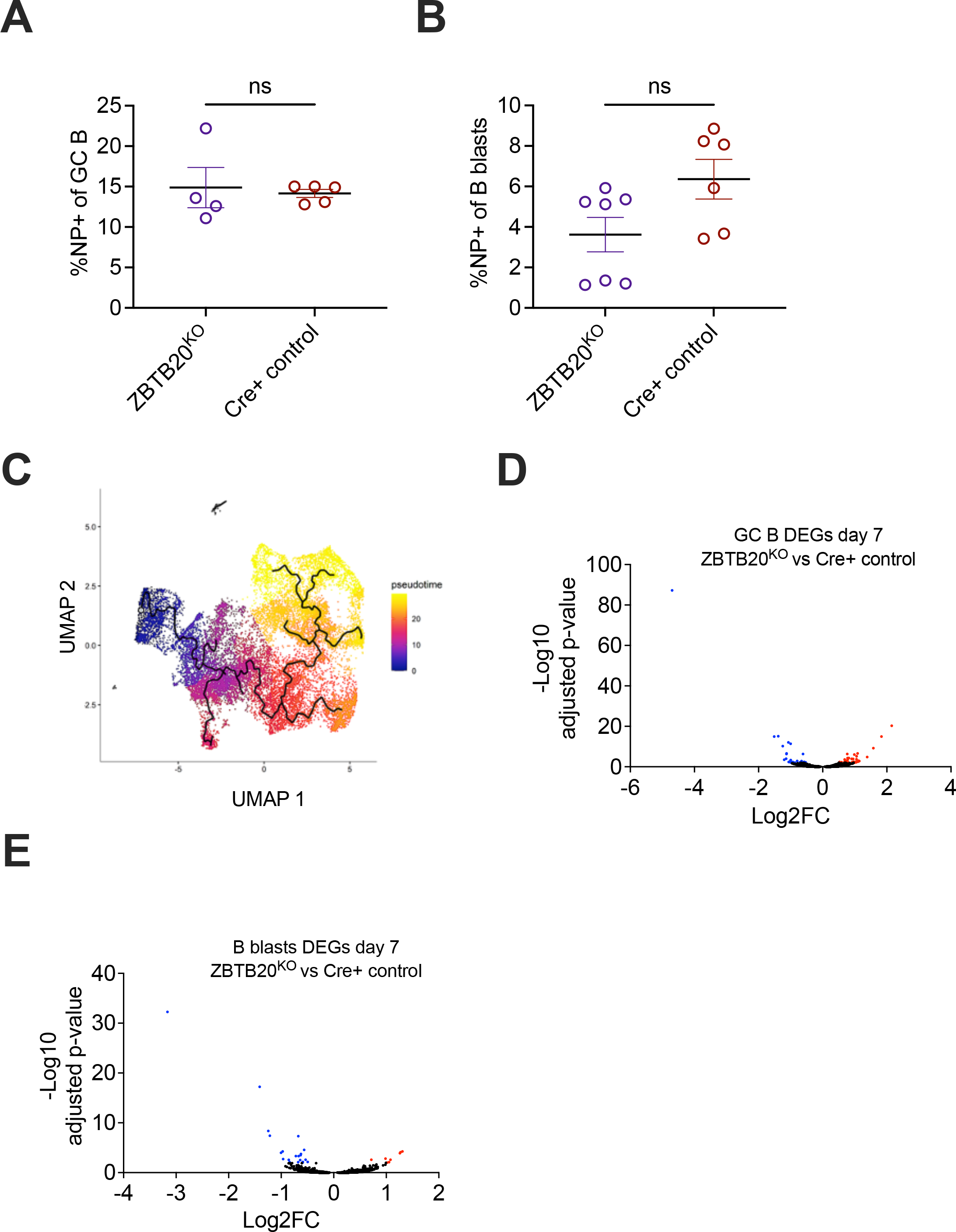
Loss of ZBTB20 does not reduce antigen-specific B cell populations and induces modest changes in the transcriptome 7 days post-<u>immunization.</u>. (**A**-**B**) Mice were immunized with NP-CGG/Alum and frequencies of (A) NP-specific GC B (A) and (B) B blasts (B) in ZBTB20^KO^ (purple) and Cre+ control (red) mice were assessed by flow cytometry. Each dot represents an individual mouse. Data are representative of three individual experiments. Mean values ± SEM are shown. No statistical significance observed by Welch’s two-tailed t-test. (**C**) Monocle trajectories were overlaid on the sc-RNAseq UMAP in Figure 3A-D depicting the transcriptional pseudotime path from day 7 B blasts to day 7 GC B cells to day 14 GC B cells. (**D**-**E**) NP-specific GC B cells and B blasts were sorted from ZBTB20^KO^ and Cre+ control mice 7 days post-immunization with NP-CGG/Alum and subjected to RNA-seq. Differentially expressed genes determined by DESeq2 (adjusted p-value < 0.01) in GC B cells (C) and B blasts (D) are depicted. DEGs upregulated in ZBTB20^KO^ mice are shown in red and downregulated genes are in blue. For GC B, n=4 for ZBTB20^KO^ and n=4 for Cre+ control. For B blasts, n=5 for ZBTB20^KO^ and n=4 for Cre+ control.

**Figure S5 related to Figure 4:**
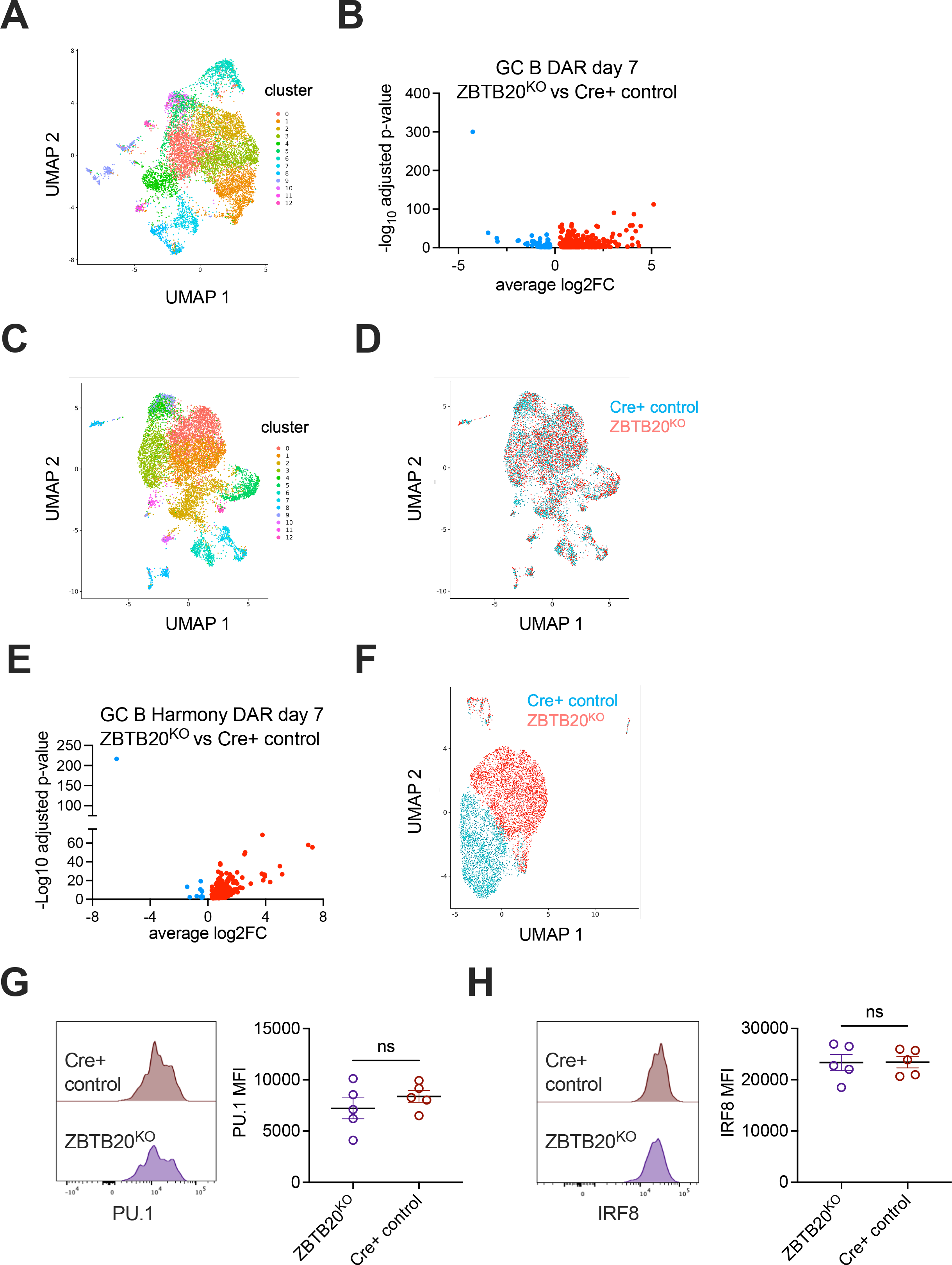
Additional analysis of epigenetic regulation and transcription factor direct repression. (**A**) UMAP showing epigenetic clustering of NP-specific ZBTB20^KO^ and Cre+ control GC B cells 7 days post immunization. Genotype identities in Figure 4A reveal enrichment of ZBTB20^KO^ cells in cluster 1 and Cre+ control cells in cluster 0. (**B**) DAR between cells in cluster 1 (ZBTB20^KO^) versus cells in cluster 0 (Cre+ control) are shown. DAR upregulated in ZBTB20^KO^ mice are shown in red and downregulated DAR are in blue. (**C**-**D**) Harmony integration was performed on single-cell epigenetic profiles from ZBTB20^KO^ and Cre+ control GC B cells 7 days post-immunization. Cluster integration showing (C) unique clusters in the total dataset, (D) successful integration of cells of indicated genotype across clusters, and (E) DARs of cells in cluster 0 are shown. (**F**) Single-cell epigenetic profiles were visualized at 21 days post-immunization in ZBTB20^KO^ (red) and Cre+ control (blue) GC B cells using Signac. Data are cumulative of cells from 7 ZBTB20^KO^ and 8 Mb1-Cre+ control mice. (**G**) PU.1 protein levels were assessed in NP specific GC B cells of ZBTB20^KO^ (purple) and Cre+ control (red) 7 days post-immunization. Representative histograms (left) and quantification (right) of PU.1 protein are indicated. Each dot represents an individual mouse. Data are representative of two individual experiments. Mean values ± SEM are shown. No statistical significance <p=0.05 was observed by an unpaired student’s two-tailed t-test. (**H**) Cells in (G) were also examined for IRF8 protein levels. Representative histograms (left) and quantification (right) of IRF8 protein are shown. Each dot represents an individual mouse. Data are representative of two individual experiments. Mean values ± SEM are shown. No significance was observed by an unpaired student’s two-tailed t-test.

**Figure S6 related to Figure 6:**
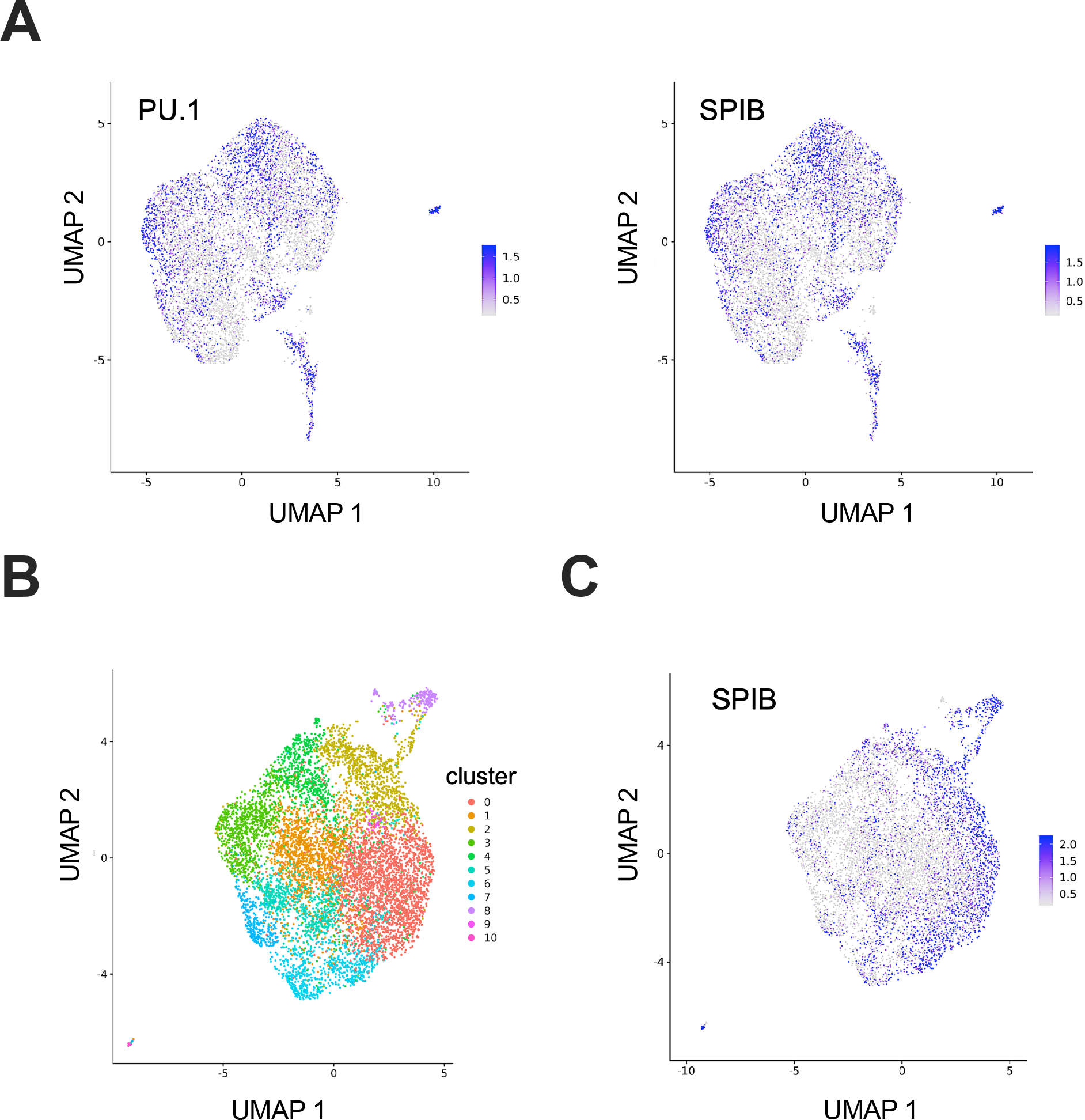
Additional epigenetic analysis of Sigma adjuvanted immunizations. (**A**) ChromVAR scoring of PU.1 (left) and SPIB (right) in GC B cells isolated 14 days post-immunization from ZBTB20^KO^ and Cre+ control mice. (**B**) Epigenetic clustering pattern of ZBTB20^KO^ GC B cells isolated 14 days post-immunization with NP-CGG adjuvanted with Alum and Sigma Adjuvant. (**C**) ChromVAR scoring of ZBTB20^KO^ Alum adjuvanted enriched cluster 2 indicates statistically significant enrichment of SPIB consensus binding motifs (adj. p-value = 2.04 x 10^-131^). scATAC-seq libraries were generated from sorted antigen-specific GC B cells. Day 14 NP-CGG/alum samples are identical to those in Figure 4. For additional day 14 NP-CGG/Sigma adjuvant libraries, data are cumulative of cells from 9 ZBTB20^KO^ and 9 Mb1-Cre+ control mice.

**Supplemental Table 1 related to Figure 3: Differential expressed genes from day 7 ZBTB20^KO^ antigen-specific GC B using scRNA-seq.** Antigen-specific cells from ZBTB20^KO^ day 7 GC B and Cre+ control day 7 GC B were assessed for differential gene expression by scRNA-seq. Antigen-specific cells from a minimum of four mice were pooled for transcriptome library construction using the 10x Genomics platform. Downstream analysis was conducted using Seurat.

**Supplemental Table 2 related to Figure 3: Differentially Expressed Genes (DEGs) from ZBTB20^KO^ day 7 antigen-specific GC B using bulk RNA-seq.** Differential gene expression analysis was conducted on antigen-specific cells using the DESeq2 extension available on the Galaxy server. Four biological replicates were used for each condition.

**Supplemental Table 3 related to Figure 3: Differentially Expressed Genes (DEGs) from ZBTB20^KO^ day 7 antigen-specific B blasts using bulk RNA-seq.** Differential gene expression analysis was conducted on antigen-specific cells using the DESeq2 extension available on the Galaxy server. 5 biological replicates were used for ZBTB20^KO^ B blasts, and 4 were used for Cre+ control.

**Supplemental Table 4 related to Figure 3: Differential expressed genes from day 14 ZBTB20^KO^ antigen-specific GC B using scRNA-seq.** Antigen-specific cells from ZBTB20^KO^ day 14 GC B and Cre+ control day 14 GC B were assessed for differential gene expression by scRNA-seq. Antigen-specific cells from a minimum of five mice were pooled for transcriptome library construction using the 10x Genomics platform. Downstream analysis was conducted using Seurat.

**Supplemental Table 5 related to Figure 4: scATAC-seq differentially accessible regions (DAR) of day 7 ZBTB20^KO^ antigen-specific GC B.** Differentially accessible regions were identified in antigen-specific cells from day 7 GC B between the ZBTB20^KO^-associated cluster 1 and the Cre+ control-associated cluster 0. Analysis was conducted using Signac.

**Supplemental Table 6 related to Figure 4: scATAC-seq DAR from Day 7 ZBTB20KO GC B with Harmony integration.** Differentially accessible regions were identified in antigen-specific cells by genotype within cluster 0 of the Harmony-integrated Signac dataset from day 7 GCB of ZBTB20^KO^ and Cre+ controls.

**Supplemental Table 7 related to Figure 4: ChromVAR results from scATAC-seq Day 7 ZBTB20^KO^ GC B.** Antigen-specific cells from day 7 GC B ZBTB20^KO^-associated clusters 1 and 3 were compared to Cre+ control-associated cluster 0 and analyzed using ChromVAR for enriched transcription factor binding motifs in ZBTB20^KO^ GC B.

**Supplemental Table 8 related to Figure 4: ChromVAR results scATAC-seq day 14 ZBTB20KO GC B.** ChromVAR was employed on antigen-specific cells from day 14 GC B ZBTB20^KO^-associated cluster 1 compared to Cre+ control-associated cluster 0 to assess transcription factor motifs with enhanced potential binding sites in ZBTB20^KO^ GC B cells.

**Supplemental Table 9 related to Figure 4: Predicted ZBTB20 binding sites in day 7 GC B DAR.** Day 7 DAR from scATAC-seq analysis were assessed for enrichment of two ZBTB20 consensus binding motif sites using the FIMO tool from the MEME suite.

**Supplemental Table 10 related to Figure 4: Overlap between predicted IRF8 and ZBTB20 binding sites within DAR.** DAR from day 7 GC B scATAC-seq analysis with predicted consensus ZBTB20 binding motif sites were assessed for enrichment of AICE, EICE, and ISRE, consensus binding motifs using the FIMO tool from the MEME suite.

**Supplemental Table 11 related to Figure 6: ChromVAR results scATAC-seq day 14 ZBTB20KO GC B Sigma adjuvant.** ChromVAR was employed on antigen-specific cells from day 14 GC B ZBTB20^KO^ associated cluster 0 compared to Cre+ control associated cluster 1 to assess transcription factor motifs with enhanced potential binding sites in ZBTB20^KO^ GC B.

**Supplemental Table 12 related to Figure 6: ChromVAR results from day 14 ZBTB20KO alum GC B enriched cluster.** ChromVAR was employed on antigen-specific cells from day 14 GC B ZBTB20^KO^ alum adjuvant associated cluster 2 compared to all other clusters to assess transcription factor motifs with enhanced potential binding sites in ZBTB20^KO^ alum adjuvant GC B.

**Supplemental Table 13 related to all Figures: Genotyping primers utilized in this study.**

